# Cross-domain encoding models reveal shared and domain-specific neural representations across language and mathematics

**DOI:** 10.64898/2026.06.07.730750

**Authors:** Tomoya Nakai, Rieko Kubo, Shinji Nishimoto

**Affiliations:** Graduate School of Information Science and Technology, the University of Tokyo, Tokyo, Japan; Center for Information and Neural Networks, National Institute of Information and Communications Technology, Suita, Japan; Graduate School of Medical and Dental Sciences, Institute of Science Tokyo, Tokyo, Japan; Graduate School of Frontier Biosciences, Osaka University, Suita, Japan; Graduate School of Medicine, Osaka University, Suita, Japan

**Keywords:** fMRI, encoding model, language, mathematics, large language model

## Abstract

Whether language and mathematics rely on shared or distinct neural representations remains an unresolved question in cognitive neuroscience. Here we combine latent features from a large language model (LLM) with vertex-wise encoding models to examine cross-domain generalization between language and mathematics. Thirty-two participants performed sentence comprehension and calculation tasks during fMRI, and encoding models were trained using features embedded in a common latent space. Cross-domain prediction identified cortical regions associated with partially shared representations, most prominently the left 55b, while control analyses suggested that these effects could not be fully explained by low-level visual processing or simple task-general factors. Task-specificity contrasts revealed stronger language-related prediction in the left anterior superior temporal and angular gyri and math-related prediction in the left precentral and intraparietal sulci. Model-weight analyses further showed that shared and domain-specific prediction patterns were reflected in distinct weight profiles across cortical regions. Connectivity analyses showed task-dependent functional coupling between cross-domain regions and language– or math-related networks. Together, these findings suggest that language and mathematics involve partially shared neural representations alongside domain-specific cortical organization, helping reconcile previous contrasting views on their neural basis.

## Introduction

A prevailing view in cognitive neuroscience posits that language and mathematics are supported by distinct neural circuits (Fedorenko, Piantadosi, & Gibson, 2024; Fedorenko & Varley, 2016; Mahowald et al., 2024). Neuroimaging studies on language have shown the involvement of the left inferior frontal and superior temporal cortices (Fedorenko et al., 2024; Friederici, Chomsky, Berwick, Moro, & Bolhuis, 2017; Price, 2012), whereas mathematical processing is more closely associated with the dorsolateral prefrontal and parietal cortices (Arsalidou, Pawliw-Levac, Sadeghi, & Pascual-Leone, 2018; Nieder, 2025; Nieder & Dehaene, 2009). This distinction has been further supported by studies that directly contrasted brain activity between language and mathematical tasks, consistently demonstrating their dissociation (Amalric & Dehaene, 2019; Fedorenko, Behr, & Kanwisher, 2011; Monti, Parsons, & Osherson, 2012). In addition, neuropsychological evidence has shown that some patients with aphasia retain relatively preserved mathematical abilities despite severe language impairments (Benn et al., 2013; Klessinger, Szczerbinski, & Varley, 2007; Varley, Klessinger, Romanowski, & Siegal, 2005). Collectively, these findings support the view that language and mathematics rely, at least in part, on distinct neural systems.

In contrast, other studies have suggested that shared neural mechanisms exist between language and mathematics. For example, some neuroimaging studies have reported overlapping activations between these two domains in the left frontal cortex (Makuuchi, Bahlmann, & Friederici, 2012; Nakai & Okanoya, 2018, 2020). Brain lesions around this region have been shown to affect both language and mathematical abilities (Baldo & Dronkers, 2007; Proios, Tsakpounidou, Karapanayiotides, Priftis, & Semenza, 2021; Semenza et al., 2006). The shared neural substrates may reflect structural properties common to both domains (Friedrich & Friederici, 2009; Hung et al., 2015; Nakai & Okanoya, 2018; Nakai & Sakai, 2014), as mathematical expressions contain structured relationships among numbers and operators analogous to those observed in natural language (Matsumoto & Nakai, 2023; Scheepers & Sturt, 2014; Scheepers et al., 2011; Schneider, Maruyama, Dehaene, & Sigman, 2012; Van de Cavey & Hartsuiker, 2016). Moreover, activity in the left frontal cortex has also been shown to reflect structural properties in geometric shapes (Al Roumi, Planton, Wang, & Dehaene, 2023; L. Wang et al., 2019), suggesting that partially overlapping cortical systems may contribute to multiple symbolic domains (Dehaene, Al Roumi, Lakretz, Planton, & Sablé-Meyer, 2022; Dehaene, Sablé-Meyer, & Ciccione, 2025). Together, these findings contrast with previously reported dissociations between language and mathematics, showing a gap between perspectives emphasizing neural independence and those supporting shared mechanisms.

Cross-domain encoding models could help bridge this gap by quantitatively evaluating feature-based brain representations and their generalizability (Dupré la Tour, Visconti di Oleggio Castello, & Gallant, 2025; Naselaris, Kay, Nishimoto, & Gallant, 2011). Encoding models, often leveraging latent features from large language models (LLMs), have been widely used to predict brain activity and to characterize neural representations of language (Caucheteux, Gramfort, & King, 2023; Deniz, Nunez-Elizalde, Huth, & Gallant, 2019; Goldstein et al., 2025; Huth, de Heer, Griffiths, Theunissen, & Gallant, 2016; Nakai, Yamaguchi, & Nishimoto, 2021; Tuckute et al., 2024) and mathematics (Nakai & Nishimoto, 2023b, 2023a). The success of these models may partly stem from the ability of LLMs to embed both linguistic and mathematical symbol sequences within a shared latent feature space (**Figure 1A**). Furthermore, by evaluating whether model predictions generalize across domains, encoding models may provide a quantitative framework for characterizing partially shared neural representations. For example, previous studies have demonstrated generalization of neural prediction across different modalities, formats, and stimulus domains, including between story listening and text reading (Nakai et al., 2021), between story listening and movie perception (Tang, Du, Vo, Lal, & Huth, 2023), and between mathematical expressions and math-word stimuli (Nakai & Nishimoto, 2023b).

**Figure 1.**
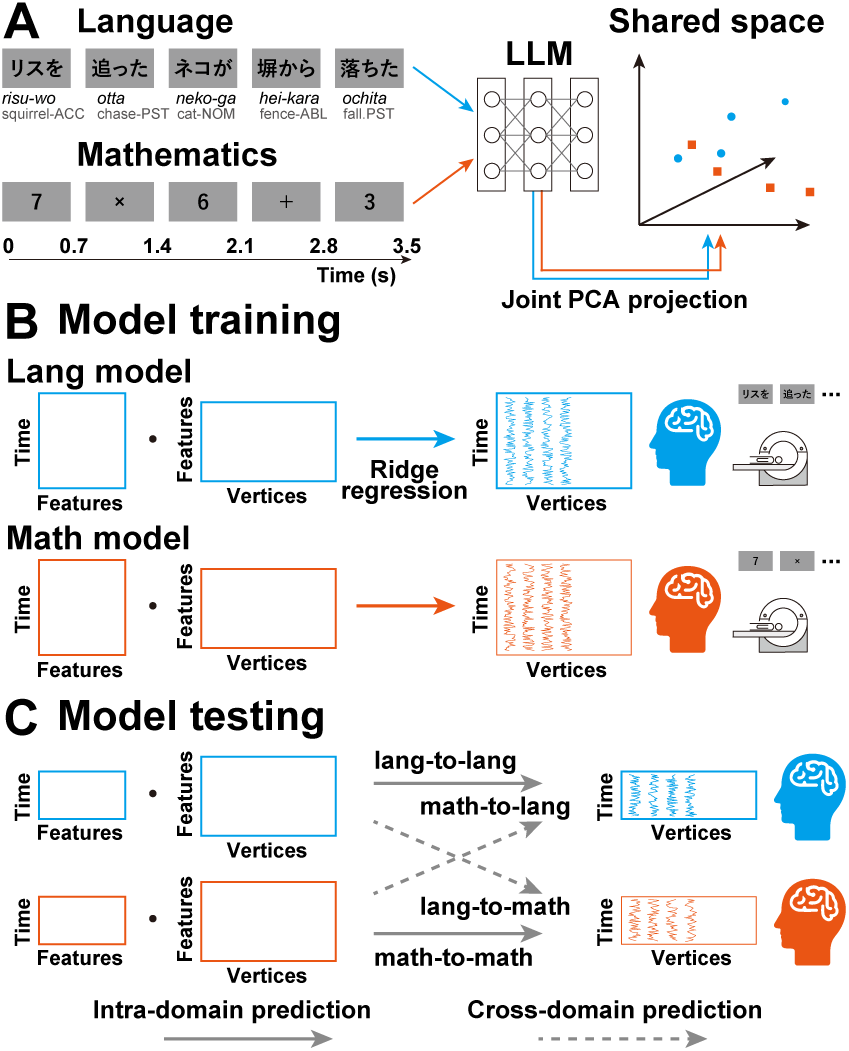
Feature extraction and encoding model analysis. **(A)** Latent features were extracted from both language (“Lang”, blue) and mathematics (“Math”, red) stimuli using a large language model (LLM), embedded into a shared feature space using principal component analysis (PCA)-based dimensionality reduction. **(B)** Vertex-wise encoding models were trained separately for each domain to predict cortical activity from the latent features (referred to as the language and math encoding models, respectively). **(C)** In the intra-domain prediction analysis (depicted by solid arrows), the language encoding model was tested on independent language test datasets (lang-to-lang), and the math encoding model was tested on independent math test datasets (math-to-math). In the cross-domain prediction analysis (depicted by dashed arrows), the language encoding model was tested on math test datasets (lang-to-math), and the math encoding model was tested on language test datasets (math-to-lang).

Here we extend this framework to characterize shared and domain-specific neural representations across language and mathematics using latent LLM features and cross-domain modeling. In this framework, the LLM served as a common feature extractor for embedding linguistic and mathematical stimuli into a shared latent space, rather than as a model of the neural computations underlying these domains. Based on previous literature (Arsalidou et al., 2018; Fedorenko et al., 2024; Friederici et al., 2017; Mahowald et al., 2024; Makuuchi et al., 2012; Nakai & Okanoya, 2018, 2020; Nieder, 2025; Nieder & Dehaene, 2009; Price, 2012), we hypothesized that cross-domain prediction would identify cortical regions showing shared representational components across language and mathematics, particularly within the left frontal cortex, while temporal and parietal regions would exhibit stronger domain specificity. To investigate this hypothesis, we conducted functional magnetic resonance imaging (fMRI) experiments with thirty-two participants who performed both sentence comprehension and calculation tasks. For each participant, we constructed vertex-wise encoding models using 100-dimensional latent features extracted from sentences and mathematical expressions by the same LLM (Llama3) (**Figure 1**). The language and mathematical stimuli were designed to have comparable symbolic structures, enabling us to evaluate whether model predictions generalized across domains. Model-weight patterns associated with shared and domain-specific components were further examined using weight correlation and principal component analysis (PCA). Finally, we tested whether the cross-domain regions exhibit distinct connectivity patterns with language– and math-related networks using functional connectivity analyses of actual and predicted brain activity. The present study contributes to reconciling contrasting views on the neural bases of language and mathematics through a unified modeling framework grounded in LLM-derived features.

## Results

### Language and math encoding models predicted activity in the left precentral and bilateral occipital cortices

We first examined whether the encoding models could predict brain activity when the training and test stimuli belonged to the same cognitive domain (i.e., *intra-domain analysis*). For each participant, we trained encoding models using 100-dimensional latent features extracted from the intermediate (16^th^) layer of the LLM (Llama3). The dimensionality-reduction step was fitted only on the training stimuli and then applied to the independent test stimuli and unstructured control stimuli, ensuring that no test information was used in feature-space construction. These models were trained either on the language stimuli (*language encoding models*) or on the math stimuli (*math encoding models*).

The language encoding models significantly predicted brain activity during the language test condition across large portions of the bilateral temporal, occipital, and frontal cortices (Wilcoxon signed-rank test, false discovery rate (FDR)-corrected *p* < 0.05; **Figure 2A**, shown in blue). Similarly, the math encoding models significantly predicted brain activity during the math test condition across extensive regions of the bilateral parietal, occipital, and frontal cortices (**Figure 2A**, shown in red). Notably, both language and math encoding models significantly predicted activity in the dorsal portion of the left precentral gyrus, corresponding to the area 55b (Glasser et al., 2016) (see **Supplementary Figure S1** for its anatomical location), as well as in the bilateral occipital cortices (**Figure 2A**, shown in white).

**Figure 2.**
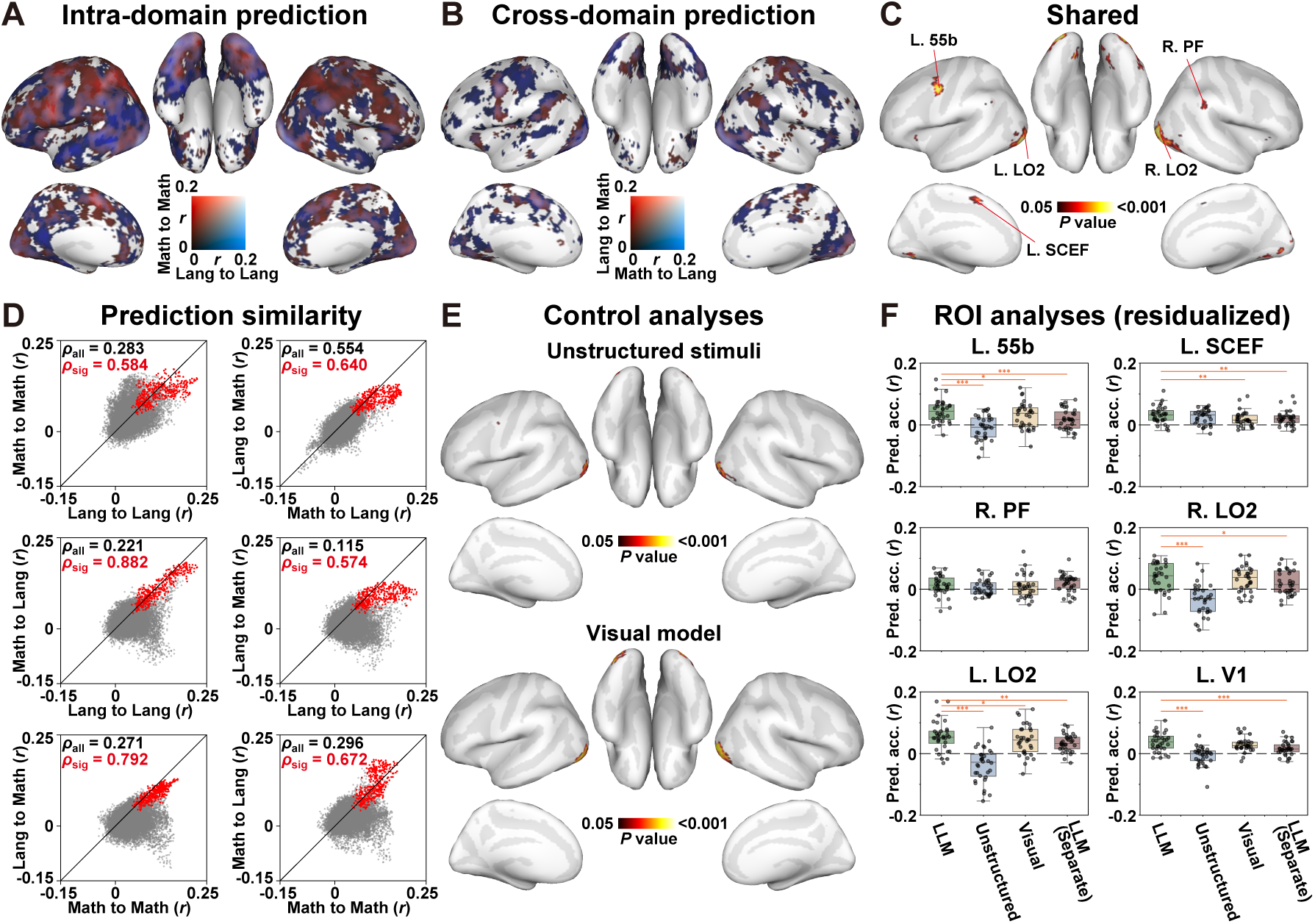
Latent LLM features predict brain activity in both language and mathematics. **(A)** Cortical maps showing brain regions with significant intra-domain prediction based on latent features from the 16th layer of the LLM (Llama3) model. The mean prediction accuracy for the language target condition using the language encoding model (*lang-to-lang*) is shown in blue, and that for the math target condition using the math encoding model (*math-to-math*) is shown in red. Overlapping regions predicted in both conditions are shown in white. **(B)** Cortical maps showing brain regions with significant cross-domain prediction. The mean prediction accuracy for the language target condition using the math encoding model (*math-to-lang*) is shown in blue, and that for the math target condition using the language encoding model (*lang-to-math*) is shown in red. Overlapping regions are shown in white. **(C)** Brain regions significant in both intra– and cross-domain predictions, revealed by conjunction analysis using FDR-corrected *p* values. **(D)** Scatter plots demonstrate correlations of prediction accuracies across all cortical vertices (colored in gray), shown for six combinations of intra-and cross-domain predictions. The subset of each plot (colored in red) indicates correlation only using significant vertices in the conjunction analysis. Correlation coefficients are computed using Spearman’s rank correlation. **(E)** Control analyses based on conjunction maps using the LLM model with unstructured stimuli and the Visual (AlexNet) model with the original test stimuli. (**F**) Box plots present mean prediction accuracy across intra– and cross-domain predictions, computed using residualized brain responses after removing task-evoked boxcar components. Results are shown for the original LLM (Llama3) model, LLM with unstructured stimuli, Visual model, and LLM with separate PCA projection. Prediction accuracies were extracted from anatomical regions-of-interest (ROIs) of left 55b, supplementary and cingulate eye field (SCEF), bilateral lateral occipital area 2 (LO2), right parietal area F (PF), and left primary visual cortex (V1). Dots represent individual participants. Asterisks indicate significance levels in ROI-based comparisons: *, *p* < 0.05; **, *p* < 0.01; ***, *p* < 0.001.

The intra-domain analysis can shed light on brain regions that contribute to both language and mathematics, confirming that our encoding models successfully predict well-established brain networks associated with these two cognitive domains. However, this analysis still leaves open the possibility that activations induced by language and mathematical tasks merely overlap in these regions, without necessarily sharing similar neural representations. To address this issue, we applied the encoding models to the test stimuli that were not used for model training (i.e., *cross-domain analysis*). When the math encoding models were tested with the language test dataset, we found significant predictions (Wilcoxon signed-rank test, FDR-corrected *p* < 0.05) in somewhat reduced yet distributed regions, including the left precentral gyrus, left superior temporal gyrus, right angular gyrus, and bilateral occipital cortices (**Figure 2B**, shown in blue). Conversely, when the language encoding models were tested with the math test dataset, we again observed significant predictions in the left precentral gyrus, right angular gyrus, and bilateral occipital cortices (**Figure 2B**, shown in red). Notably, both the language and math encoding models significantly predicted activity in the left 55b and bilateral occipital cortices (**Figure 2B**, shown in white).

To identify brain regions showing cross-domain generalization across the two domains, we performed a conjunction analysis between intra– and cross-domain predictions. Based on the minimum-statistic approach (Nichols, Brett, Andersson, Wager, & Poline, 2005), each vertex was assigned the smallest prediction accuracy among the two intra-domain (lang-to-lang, math-to-math) and two cross-domain (lang-to-math, math-to-lang) analyses. The results revealed significant predictions in the left 55b, left middle cingulate cortex (SCEF), right inferior parietal cortex (PF), and bilateral lateral occipital cortices including the LO2 (**Figure 2C, Table S1**). Similar regions were identified using a block-permutation test (**Supplementary Figure S2**). These regions were hereafter referred to as *cross-domain regions*.

Spatial prediction patterns were highly similar between language and math encoding models (**Figure 2D**). Prediction accuracies were positively correlated for all combinations (Lang-to-Lang and Math-to-Math, Spearman’s correlation coefficient, *ρ* = 0.283; Math-to-Lang and Lang-to-Math, *ρ* = 0.554; Lang-to-Lang and Math-to-Lang, *ρ* = 0.221; Lang-to-Lang and Lang-to-Math, *ρ* = 0.115; Math-to-Math and Lang-to-Math, *ρ* = 0.271; Math-to-Math and Math-to-Lang, *ρ* = 0.296), and the correlation became larger when using only significant vertices in the conjunction analysis (Lang-to-Lang and Math-to-Math, *ρ* = 0.584; Math-to-Lang and Lang-to-Math, *ρ* = 0.640; Lang-to-Lang and Math-to-Lang, *ρ* = 0.882; Lang-to-Lang and Lang-to-Math, *ρ* = 0.574; Math-to-Math and Lang-to-Math, *ρ* = 0.792; Math-to-Math and Math-to-Lang, *ρ* = 0.672). These findings suggest partially shared neural representations across language and mathematics.

### Cross-domain prediction extends beyond simple visual or memory factors and is reproducible across different LLMs

Sentence comprehension and calculation tasks may involve various cognitive factors, including visual processing, memory retrieval, and spatial attention. The significant predictions observed in the left 55b, SCEF, right PF, and bilateral lateral occipital cortices by the encoding models might be explained by these general cognitive factors, or by methodological factors such as feature dimensionality, model capacity, or properties of LLM-derived latent features unrelated to shared representations. To address these possibilities, we performed three control analyses testing the effect of (i) stimulus datasets and task demands, (ii) low-level visual features and model dimensionality, and (iii) feature-space construction, respectively.

First, to evaluate the influence of stimulus datasets and task demands, we applied the same language and math encoding models to different test datasets (i.e., the unstructured stimuli in the control condition). In the control run, the elements within each linguistic or mathematical stimulus could not be integrated into a coherent sentence or calculable expression (e.g., five unrelated verbs or five digits), thereby minimizing structural relationships among stimulus elements (**Supplementary Figure S3A**). Importantly, however, the control stimuli matched the test stimuli in the number of presented items, and participants were required to memorize these items. Thus, the demands on low-level visual processing, short-term memory, and attention were comparable between the control and test stimuli. The whole-brain conjunction analysis showed limited prediction accuracy in the left 55b with the control stimuli, while prediction patterns were relatively preserved in the bilateral lateral occipital cortices (**Figure 2E** top).

To examine the differences in predictive performance between conditions in greater detail, we extracted mean prediction accuracy in the predetermined anatomical ROIs. These ROIs included anatomical regions corresponding to the cross-domain regions identified in the conjunction analysis (left 55b, left SCEF, right PF, left LO2, and right LO2), as well as the left V1, which was included to further assess the contribution of early visual processing. We then compared prediction accuracy between the original test stimuli and the control stimuli. Prediction accuracy was higher with the original test stimuli in the left 55b (Wilcoxon signed-rank test, *p* = 0.039, rank-biserial correlation, *r_rb_* = 0.242; **Supplementary Figure S4**), left SCEF (*p* = 0.021, *r_rb_* = 0.271), right PF (*p* = 0.007, *r_rb_* = 0.328), and left V1 (*p* < 0.001, *r_rb_* = 0.471), but not in the left LO2 (*p* = 0.053, *r_rb_* = 0.232) or right LO2 (*p* = 0.185, *r_rb_* = 0.182).

We further repeated this ROI analysis using residualized brain responses after removing task-evoked boxcar components (see **Supplementary Information**). After residualization, prediction accuracy remained higher with the original test stimuli than with the control stimuli, with a more pronounced difference in the left 55b (*p* < 0.001, *r_rb_* = 0.684), left LO2 (*p* < 0.001, *r_rb_* = 0.818), right LO2 (*p* < 0.001, *r_rb_* = 0.689), and left V1 (*p* < 0.001, *r_rb_* = 0.725), but not in the left SCEF (*p* = 0.287, *r_rb_* = 0.094) or right PF (*p* = 0.102, *r_rb_* = 0.197) (**Figure 2F**). Together, these analyses suggest that cross-domain prediction in the left 55b was not fully explained by simple task-general factors or mean task-evoked responses, whereas prediction in the left LO2 may partly reflect stimulus-format or visual-processing factors common to both conditions.

Second, to evaluate the influence of visual processing, we trained an alternative encoding model (*Visual* model) based on the latent visual features and calculated prediction performance with the original test stimuli (**Supplementary Figure S3B**). Specifically, 100-dimensional latent features were extracted from stimulus images using AlexNet, a convolutional neural network widely used to model visual processing in the human brain (Gu, Jamison, Sabuncu, & Kuceyeski, 2022; A. Y. Wang, Kay, Naselaris, Tarr, & Wehbe, 2023; Wen, Shi, Chen, & Liu, 2018), and language and math encoding models were tested in the same way as the LLM-based models. We expected that the Visual model would predominantly capture the contribution of low-level visual information in the current experiment. In addition, this model was based on the shared latent features projected onto the same 100-dimensional space, as was done for the LLM-based models, controlling for potential effects of dimensionality of the latent variables (see also **Supplementary Figure S5** for the LLM-based model performance with different numbers of dimensions). Using the conjunction analysis, only bilateral lateral occipital cortices were significantly predicted (**Figure 2E** bottom). ROI analyses further revealed higher prediction accuracy of the LLM-based model relative to the Visual model in the original test run in the left 55b (*p* < 0.001, *r_rb_* = 0.340; **Supplementary Figure S4**), left SCEF (*p* < 0.001, *r_rb_* = 0.260), left LO2 (*p* < 0.001, *r_rb_* = 0.301), and right LO2 (*p* = 0.008, *r_rb_* = 0.186), but not in the right PF (*p* = 0.053, *r_rb_* = 0.152) or left V1 (*p* = 0.211, *r_rb_* = 0.080).

This pattern was also observed when residualized brain responses were used: the LLM-based model showed higher prediction accuracy than the Visual model in the left 55b (*p* = 0.012, *r_rb_* = 0.213), left SCEF (*p* = 0.003, *r_rb_* = 0.326), and left LO2 (*p* = 0.045, *r_rb_* = 0.139), but not in the right PF (*p* = 0.373, *r_rb_* = 0.104), right LO2 (*p* = 0.112, *r_rb_* = 0.103), or left V1 (*p* = 0.075, *r_rb_* = 0.162) (**Figure 2F**). These findings suggest that the prediction patterns observed in the language and math encoding models cannot be explained solely by low-level visual processing or by the use of 100-dimensional feature vectors.

Third, to check the importance of our feature space construction (i.e., joint PCA projection), we trained encoding models using the same LLM (Llama3) and the same test datasets, but with a different feature preprocessing pipeline. Specifically, we projected LLM features onto separate 100-dimensional spaces for language and math stimuli (i.e., separate PCA projection; **Supplementary Figure S3C**), and tested intra– and cross-domain prediction performance. No brain region showed significant prediction in the conjunction analysis. ROI analyses further revealed higher prediction accuracy for the joint PCA model than for the separate PCA model in the left 55b (*p* < 0.001, *r_rb_* = 0.447; **Supplementary Figure S4**), left SCEF (*p* < 0.001, *r_rb_* = 0.333), left LO2 (*p* < 0.001, *r_rb_* = 0.574), and right LO2 (*p* < 0.001, *r_rb_* = 0.541), but not in the right PF (*p* = 0.345, *r_rb_* = 0.008) or left V1 (*p* = 0.053, *r_rb_* = 0.195).

Using residualized brain responses, the joint PCA model still showed higher prediction accuracy than the separate PCA model in the left 55b (*p* < 0.001, *r_rb_* = 0.381), left SCEF (*p* = 0.006, *r_rb_* = 0.240), left LO2 (*p* = 0.003, *r_rb_* = 0.301), right LO2 (*p* = 0.023, *r_rb_* = 0.254), and left V1 (*p* < 0.001, *r_rb_* = 0.445) but not in the right PF (*p* = 0.815, *r_rb_* = –0.135) (**Figure 2F**). These results indicate that cross-domain prediction critically depends on mapping language and mathematical stimuli into a shared latent space. Thus, the observed cross-domain prediction cannot be attributed simply to the use of 100-dimensional LLM-derived features, but requires a common projection space that preserves correspondences between the two domains.

As an additional control, we tested whether cross-domain prediction depended on the correct temporal correspondence between stimuli and LLM-derived features. We circularly shifted the training feature time series within each run and repeated the encoding analysis 100 times (**Supplementary Information**). The mean prediction map across the 100 shifted-feature models showed no significant regions in the conjunction analysis. This result suggests that cross-domain prediction was not driven solely by temporal structure in the BOLD responses shared across the language and math tasks, but depended on the correct temporal correspondence between stimuli and LLM-derived features.

Having addressed potential confounds related to general cognitive factors and feature preprocessing, we next examined whether distinct transformer subcomponents contributed differently to cross-domain and domain-specific prediction. Specifically, we compared representations derived from the self-attention and MLP sublayers of the 16th transformer block. Attention– and MLP-derived features exhibited distinct spatial patterns of prediction (**Supplementary Figure S6**). Attention-derived features tended to show significant cross-domain prediction across broader regions of the cortex, whereas MLP-derived features were more associated with domain-specific prediction. This pattern suggests a difference in the extent to which the two sublayers contribute to cross-domain versus domain-specific representations.

To further examine reproducibility and model– and layer-dependency of our findings, we extracted latent features from five distinct layers of Llama3 (1^st^, 8^th^, 16^th^, 24^th^, and 32^nd^ layers), Gemma2 (1^st^, 7^th^, 13^th^, 20^th^, and 26^th^ layers), Qwen2.5 (1^st^, 7^th^, 14^th^, 21^st^, and 28^th^ layers), GPT-NeoX (1^st^, 8^th^, 16^th^, 24^th^, and 32^nd^ layers), RakutenAI2 (1^st^, 8^th^, 16^th^, 24^th^, and 32^nd^ layers), and LLM-jp (1^st^, 10^th^, 20^th^, 30^th^, and 40^th^ layers), and constructed lang– and math-encoding models. We found prediction patterns similar to those of the main Llama3 16th-layer model, in that conjunction analysis showed significant prediction in the left 55b, bilateral insula, and bilateral lateral occipital cortices (**Supplementary Figure S7**).

Prediction accuracy was also comparable across different LLMs and layers (Llama3, Kruskal-Wallis test, *p* = 0.973; GPT-NeoX, *p* = 0.359; Gemma2, *p* = 0.968; Qwen2.5, *p* = 0.826; RakutenAI2, *p* = 0.907; LLM-jp, *p* = 0.938; **Supplementary Figure S8**). These results suggest that the main prediction patterns were not specific to a particular LLM or layer, but were broadly reproducible across the tested models and layers.

### Domain-specific brain regions for language and mathematics

Although the previous analyses revealed cross-domain brain regions predictable in both language and mathematics, it remained unclear whether the encoding model also captured domain-specific neural representations. To address this question, we quantified domain-specific brain regions by comparing intra-domain and cross-domain prediction accuracies. Specifically, language-specificity was quantified by contrasting the intra-domain prediction in language (*lang-to-lang*) and the average of cross-domain predictions (*lang-to-math* and *math-to-lang*) (**Figure 3A**), and math-specificity was quantified by contrasting intra-domain prediction in mathematics (*math-to-math*) and average of cross-domain predictions (*math-to-lang* and *lang-to-math*) (**Figure 3B**). We expected that the cross-domain predictions would reflect brain representations shared between language and mathematics, thereby serving as an effective control to exclude contributions from shared components present in the *lang-to-lang* and *math-to-math* analyses (i.e., intra-domain predictions).

**Figure 3.**
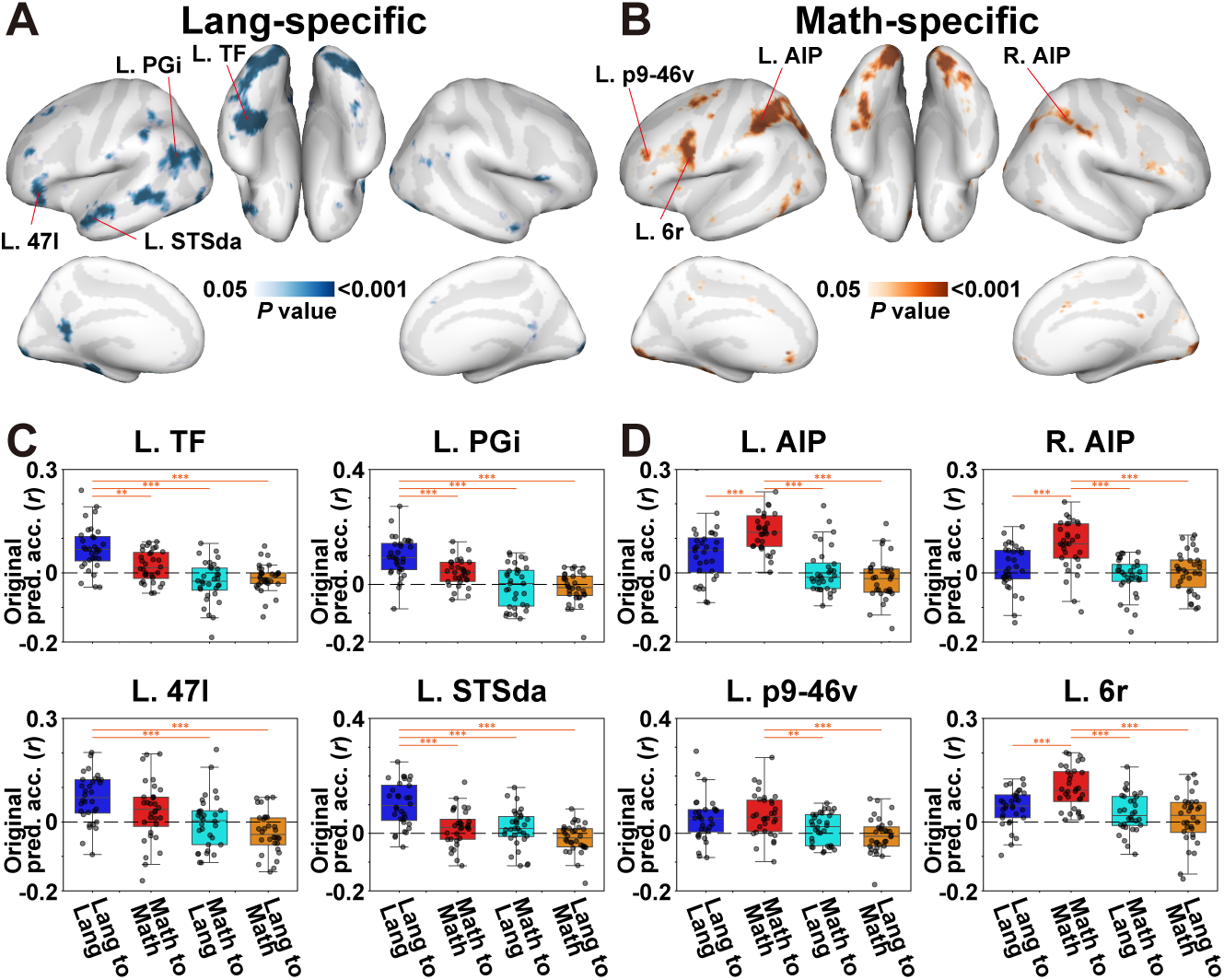
Latent LLM features predict domain-specific components of language and mathematics. **(A)** Cortical maps showing brain regions with significantly higher intra-domain prediction (*lang-to-lang*) than cross-domain prediction (average of *math-to-lang and lang-to-math*) for the language target condition, thresholded at FDR-corrected *p* < 0.05. **(B)** Cortical maps showing brain regions with significantly higher intra-domain prediction (*math-to-math*) than cross-domain prediction (average of *math-to-lang and lang-to-math*) for the math target condition. **(C-D)** Box plots demonstrate prediction accuracies of all four prediction types (*lang-to-lang*, *math-to-math*, *math-to-lang*, and *lang-to-math*), averaged across vertices within each of the target **(C)** language-specific ROIs (left TF, PGi, 47l, and STSda) and **(D)** math-specific ROIs (left AIP, right AIP, left p9-46v, and 6r). Dots represent data from each participant. Asterisks indicate significance levels in ROI-based comparisons: *, *p* < 0.05; **, *p* < 0.01; ***, *p* < 0.001.

In the language-specificity contrast, we observed significant effects in the left inferior frontal gyrus (47l), left anterior superior temporal sulcus (superior temporal sulcus dorsal anterior, STSda), the left angular gyrus (parietal area G inferior, PGi), and left fusiform gyrus (temporal fusiform, TF) (Wilcoxon signed-rank test, FDR-corrected *p* < 0.05; **Figure 3A**). Similar spatial patterns were obtained when language-specificity was defined using each cross-domain prediction separately rather than their average (**Supplementary Figure S9A-B**). Detailed investigation of prediction accuracies in these ROIs revealed that these regions showed higher intra-domain prediction in language (*lang-to-lang*) than the other prediction types (left TF, *p* < 0.002 and *r_rb_* > 0.477 for all combinations; left PGi, *p* < 0.001 and *r_rb_* = 0.529 for all combinations; left STSda, *p* < 0.001 and *r_rb_* > 0.564 for all combinations), except for the left 47l (*p* < 0.001 and *r_rb_* > 0.521 for *lang-to-lang* vs. *lang-to-math* and *math-to-lang*; *p* = 0.064 and *r_rb_* = 0.271 for *lang-to-lang* vs. *math-to-math*), suggesting that activity in these ROIs can be specifically predicted when both training and test stimuli are in the language domain.

In the math-specificity contrast, significant effects were found in the left precentral sulcus (rostral area 6, 6r), left posterior lateral prefrontal cortex (p9-46v), and the bilateral intraparietal sulcus (anterior intraparietal, AIP) (**Figure 3B**). Similar spatial patterns were obtained when math-specificity was defined using each cross-domain prediction separately rather than their average (**Supplementary Figure S9C-D**). These ROIs showed higher intra-domain prediction in mathematics (*math-to-math*) than all the other prediction types in the left AIP (*p* < 0.001 and *r_rb_* > 0.533 for all combinations), right AIP (*p* < 0.001 and *r_rb_* > 0.529 for all combinations) and left 6r (*p* < 0.001 and *r_rb_* > 0.498 for all combinations), with partially consistent patterns observed in the left p9-46v (*p* = 0.180 and *r_rb_* = 0.111 for *math-to-math* vs. *lang-to-lang*; *p* = 0.003 and *r_rb_* = 0.353 for *math-to-math* vs. *math-to-lang*; *p* < 0.001 and *r_rb_* = 0.549 for *math-to-math* vs. *lang-to-math*), suggesting that activity in these ROIs can be specifically predicted when both training and test stimuli are in the math domain. These findings are consistent with previous neuroimaging studies on language (Fedorenko et al., 2024; Friederici et al., 2017; Price, 2012) and mathematics (Arsalidou et al., 2018; Nieder, 2025; Nieder & Dehaene, 2009), suggesting that the encoding models successfully captured not only shared components but also domain-specific patterns of brain activity for both language and mathematics.

### Model-weight organization across cross-domain and domain-specific regions

To further characterize the representational organization of cross-domain and domain-specific regions captured by the encoding models, we performed two types of analyses based on the encoding model weights. First, we directly compared language and math model weights using Spearman’s correlation at each vertex and ROI. The vertex-wise weight correlation showed spatially distributed similarities and differences between language and math model weights (**Figure 4A**). Importantly, brain regions in which we found significant conjunction prediction (i.e., left 55b, left SCEF, right PF, and left LO2, with the right LO2 omitted because it showed a similar pattern to the left LO2) also exhibited positive weight correlations (**Figure 4B, Supplementary Figure S10A**), consistent with the cross-domain prediction results, suggesting partially similar feature-to-response mappings across domains in these regions. In contrast, brain regions that were specifically predicted in language or in mathematics (left TF, left PGi, left 47l, left 6r, p9-46v, and bilateral AIP, except for the left STSda) showed negative weight correlations (**Figure 4C, Supplementary Figure S10B, C**), indicating divergent feature-to-response mappings between language and mathematics.

**Figure 4.**
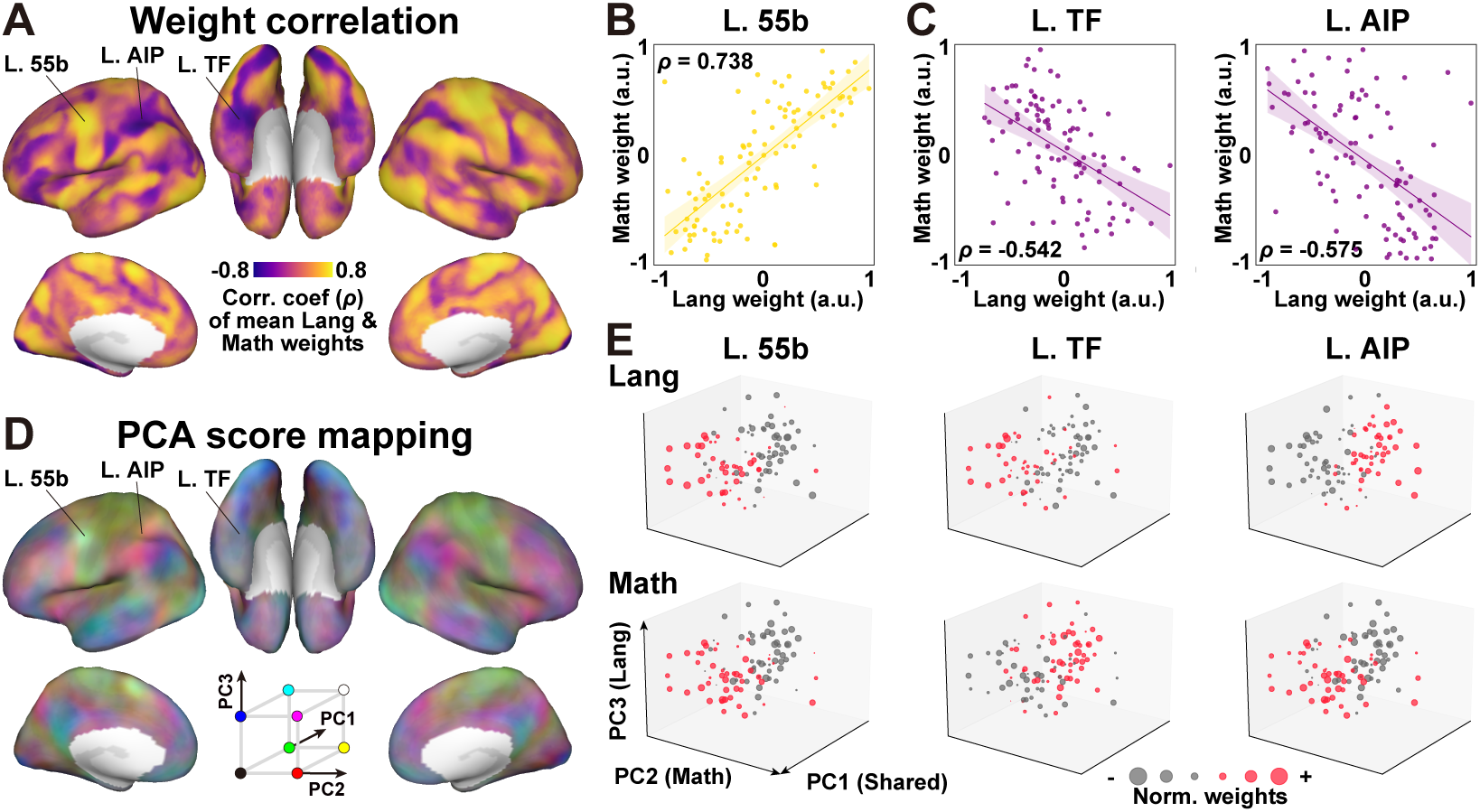
Distribution of shared and domain-specific model-weight patterns. **(A)** Weight correlation map indicates vertex-wise Spearman’s correlation between language and math model weights, averaged across participants. Vertices with a positive correlation are shown in yellow, while those with negative correlation are shown in purple. **(B-C)** Scatter plots showing correlation between language and math model weights in **(B)** the left 55b and **(C)** left TF and AIP. **(D)** Cortical map showing the distribution of the normalized scores of the top three principal components (PCs) (PC1, green; PC2, red; PC3, blue), calculated from the encoding model weights of the 1st LLM layer. **(E)** Scatter plots showing the weight values of the 100 latent features in the left 55b, TF and AIP, mapped onto a three-dimensional space defined by the PC1, PC2, and PC3 axes. Circle size represents the absolute weight value, and features with positive weights are colored in red.

Second, to visualize how the latent feature dimensions were organized in the encoding model space, we performed principal component analysis (PCA) on the model weights and mapped the first three principal components (PC1–PC3) onto the cortical surface (colored in green, red, and blue, respectively; **Figure 4D**). For each PC, we examined its spatial association with the conjunction map (**Figure 2C**), language-specificity contrast (**Figure 3A**), and math-specificity contrast (**Figure 3B**). PC1 (green) was primarily distributed across the bilateral occipital and inferior temporal cortices and showed the strongest correlation with the conjunction map (**Supplementary Figure S11B**, left), suggesting that this axis captured variance related to the cross-domain regions. PC2 (red) was distributed in the left frontal and parietal cortices and exhibited the strongest correlation with the math-specificity and conjunction maps (**Supplementary Figure S11B**, center), suggesting that this axis was aligned with math-specific prediction patterns. PC3 (blue) was distributed across the bilateral superior temporal cortices, central sulcus, and cingulate cortex and showed comparable correlations with both the language-specificity and conjunction maps (**Supplementary Figure S11B**, right), suggesting that this axis was aligned with language-specific prediction patterns.

Based on these spatial associations, we visualized how the latent feature dimensions were arranged in the space defined by the first three weight-derived PCs. Specifically, each feature dimension was represented in a three-dimensional coordinate system defined by PC1, PC2, and PC3, which were spatially associated with the conjunction, math-specificity, and language-specificity maps, respectively (**Figure 4E**). This visualization indicates that the latent feature dimensions showed preferential associations with cross-domain, language-specific, and math-specific prediction patterns. We then examined how these feature dimensions were weighted within representative ROIs: the left 55b, TF, and AIP. In each ROI, feature dimensions with positive weight values were visualized as red circles (**Figure 4E**). In the left 55b, feature dimensions located toward the left anterior side of the three-dimensional map, corresponding to the PC1 direction associated with cross-domain prediction, exhibited positive weights in both language and math encoding models (**Figure 4E** left), in line with partially shared representations across language and mathematics. In contrast, in the left TF, feature dimensions at the left anterior side had positive weights in the language encoding model but not in the math encoding model (**Figure 4E** center), consistent with the language-specific prediction pattern observed in this region (**Figure 3**). In the left AIP, feature dimensions located toward the left anterior side had positive weights most strongly in the math encoding model but not in the language encoding model, consistent with the math-specific prediction pattern observed in this region (**Figure 3**). Together, these visualizations were consistent with the prediction-based analyses, indicating that cross-domain and domain-specific regions showed distinct model-weight profiles.

Visualizations in the other ROIs demonstrated mostly similar patterns (**Supplementary Figure S12**). In the left SCEF and LO2, feature dimensions on the left anterior side exhibited positive weights in both domains. In the left PGi and 47l, these feature dimensions showed positive weights in the language encoding model but negative weights in the math encoding model. In the right AIP, left p9-46v, and left 6r, they showed positive weights in the math encoding model but negative weights in the language encoding model. In contrast, the right PF did not show the same feature-weight profile as the left 55b, and the left STSda showed a pattern distinct from the other language-specific ROIs. These findings suggest that even within cross-domain and domain-specific regions, model-weight profiles varied across ROIs.

### The left 55b and other cross-domain regions exhibit task-dependent connectivity with language– and math-related networks

Based on the observation in the domain specificity analysis and PCA, we expected that the cross-domain regions (left 55b, SCEF, PF, and LO2) would exhibit task-dependent connectivity with language– and math-specific networks. To test this possibility, we computed functional connectivity separately for the language and math tasks, using anatomically defined seed ROIs corresponding to the cross-domain regions.

We observed significantly stronger connectivity during the language task than in the math task within the bilateral inferior frontal cortex, superior and inferior temporal cortices, and along the central sulcus (Wilcoxon signed-rank test, FDR-corrected *p* < 0.05; **Figure 5A**, shown in red). Conversely, stronger connectivity during the math task was found within the bilateral lateral prefrontal, parietal, and precentral cortices (**Figure 5A**, shown in blue). A similar connectivity pattern was found in the ROI-based model connectivity analysis (**Figure 5B**), in that the left 55b seed showed stronger connectivity with the left superior temporal, left inferior temporal, and left inferior frontal ROIs during the language task, whereas the same seed showed stronger connectivity with the bilateral parietal and lateral prefrontal ROIs during the math task. The other seed ROIs (cross-domain regions: left SCEF, right PF, and left LO2) showed similar connectivity patterns (**Supplementary Figure S13A**), except for the right PF exhibiting an opposite connectivity pattern.

**Figure 5.**
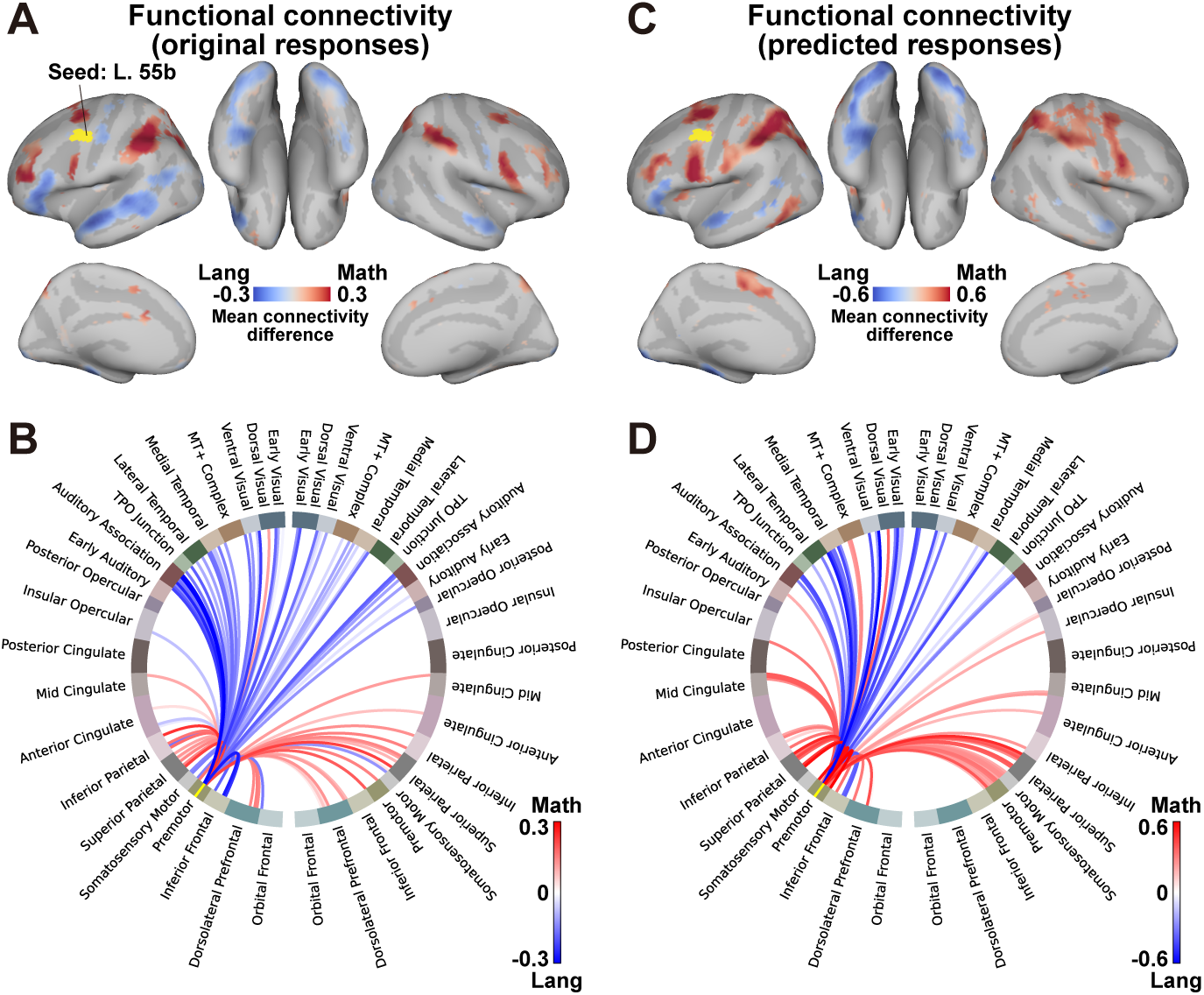
Domain-specific connectivity patterns from the left 55b. **(A)** A cortical map of vertex-wise functional connectivity from the anatomical left 55b ROI (in yellow), computed based on the original brain responses. Vertices with significantly higher connectivity during the language task are shown in red, and those with significantly higher connectivity during the math task are shown in blue (two-sided Wilcoxon signed-rank test, FDR-corrected *p* < 0.05). **(B)** A circle diagram of functional connectivity computed based on the ROI-averaged responses across 360 HCP-MMP1 ROIs (Glasser et al., 2016), summarized based on 42 different region labels and node colors. ROIs with significantly higher connectivity in language are shown in red, and those with significantly higher connectivity in mathematics are shown in blue. (**C**) A cortical map of vertex-wise functional connectivity, computed based on the predicted responses by intra-domain encoding models. (**D**) A circle diagram of functional connectivity, computed based on the ROI-averaged predicted responses by the intra-domain encoding models.

To further test whether seed-target associations were modulated by task domain, while separating this effect from overall task-domain differences, we performed a seed-by-task interaction analysis in which task-domain main effects were included as terms separate from the seed-by-task interaction term (see **Supplementary Information**). This analysis revealed a similar task-dependent pattern, with stronger left 55b associations with temporal and inferior frontal regions during the language task and stronger associations with parietal and lateral prefrontal regions during the math task (**Supplementary Figure S14**). Thus, the task-dependent association pattern was not limited to condition-wise correlations, but was also observed after separating overall task-evoked coactivation from seed-by-task modulation.

These functional connectivity results, however, do not necessarily indicate that these seed regions interact with other regions based on information captured by the latent LLM features. They might instead reflect coactivation driven by general task-related factors unrelated to the feature-based representations captured by the encoding models. To address this possibility, we further computed connectivity using the model-predicted responses generated by the intra-domain encoding models. This predicted-response connectivity analysis revealed a broadly similar task-dependent pattern (**Figure 5C, D**). For the left 55b seed, predicted responses showed stronger connectivity with the left superior temporal, left inferior temporal, and left inferior frontal ROIs during the language task, whereas stronger connectivity with bilateral parietal and lateral prefrontal ROIs was observed during the math task. Similar patterns were observed for the left SCEF and left LO2 seeds, whereas the right PF showed a divergent pattern in the predicted-response analysis (**Supplementary Figure S13B**). These findings suggest that the encoding models captured part of the task-dependent functional coupling between cross-domain regions and domain-specific networks.

## Discussion

The present study aimed to address the gap in previous literature that has reported both shared and distinct neural representations across language and mathematics, by leveraging latent LLM features and encoding models. We identified cortical regions showing cross-domain generalization across the two domains, most consistently in the left 55b, together with lateral occipital and parietal regions that may reflect partially distinct contributions. Model weight analyses revealed distinct representations within these networks, suggesting coexistence of shared and domain-specific cortical organization. Furthermore, some cross-domain regions, particularly the left 55b, exhibited task-dependent functional coupling with language– and math-related networks during their respective tasks. Together, these findings help reconcile previous contrasting views on shared and distinct neural representations across language and mathematics.

A key contribution of the present study is that language and mathematical stimuli were embedded into a shared latent feature space derived from LLMs, enabling quantitative evaluation of cross-domain prediction and shared representations. This approach made it possible to predict brain activity across language and mathematics using encoding models trained in a different domain. Previous studies have investigated the overlap of brain activity between language and mathematical tasks (Makuuchi et al., 2012; Nakai & Okanoya, 2020), examined lesions in patients with aphasia and acalculia (Baldo & Dronkers, 2007), or explored cross-domain interactions through structural priming (Nakai & Okanoya, 2018). However, these studies did not incorporate computational models capable of quantitatively evaluating neural representations, and therefore could not determine whether specific brain regions exhibit partially shared representations across language and mathematics, nor could they quantitatively assess the generalization across the two domains. The present approach helps address these limitations and provides a quantitative framework for evaluating partially shared representations across language and mathematics.

It should be noted, however, that our primary aim was to quantitatively characterize the commonalities and distinctions between language and mathematics in the brain, rather than to claim that these domains are represented identically within the LLM. While our results suggest a partial alignment between neural activity and LLM-derived representations, consistent with recent studies reporting parallels between biological and artificial neural networks (Caucheteux et al., 2023; Goldstein et al., 2022; Schrimpf et al., 2021), the internal representations within the LLM do not necessarily align completely with the neural representations underlying language and mathematics. In the current study, we used the LLM purely as a feature extractor, and the correspondence between the brain and the model was not the primary focus of our investigation. Although the LLMs used in the current study (Llama3, Gemma2, Qwen2.5, GPT-NeoX, RakutenAI2, and LLM-jp) sufficiently captured latent structure shared across language and mathematics, they are not necessarily the optimal models. Therefore, a recent report of distinct representations of language and mathematics in LLMs (Kisako, Kuribayashi, & Sasano, 2025) does not contradict our findings. Even if the shared components of language and mathematics are relatively limited within the LLM, isolating and leveraging these components as latent features enabled successful cross-domain prediction of brain activity.

The left 55b, in which cross-domain prediction was observed for both language and mathematics in the present study, was first reported in the 1950s in relation to language processing but was largely overlooked until it was revisited by the Human Connectome Project (Glasser et al., 2016; Hopf, 1956). The left 55b is located in the posterior portion of the middle frontal gyrus, inferior to the frontal eye field and superior to the ventral premotor cortex. Anatomically, it is characterized by a lower myelin content compared to the surrounding cortical regions. It has been reported to show higher activation than adjacent areas during sentence comprehension tasks and to be functionally connected to other language-related regions such as the IFG (Glasser et al., 2016). Subsequent studies have shown the involvement of this region in speech production (Chang et al., 2020; Hazem et al., 2021) and in sensorimotor integration during music perception (Siman-Tov et al., 2022; Zalta, Large, Schön, & Morillon, 2024), suggesting that this region contributes to multiple cognitive functions across domains. Regarding mathematical cognition, previous meta-analyses have demonstrated involvement of the left middle frontal gyrus across multiple types of mathematical operations (Arsalidou & Taylor, 2011; Istomina & Arsalidou, 2024). Although these studies were based on volume-based analyses, their findings are consistent with the possibility that the left 55b contributes to neural representations shared across language and mathematics. In contrast, previous studies investigating shared processing between language and mathematics have reported the involvement of the left IFG (Area 44) (Makuuchi et al., 2012; Nakai & Okanoya, 2018, 2020). This discrepancy in the reported cortical loci may stem from the fact that most previous studies employed volume-based analyses, which overlook sulcal folding patterns. In the present study, we constructed vertex-wise (i.e., surface-based) encoding models to examine how language-and math-related representations are organized across gyri and sulci. Moreover, surface-based analysis allowed us to avoid the spatial smoothing that can lead to signal contamination, thereby making our encoding models more sensitive to local activity patterns (Oosterhof, Wiestler, Downing, & Diedrichsen, 2011; Tucholka, Fritsch, Poline, & Thirion, 2012).

In addition to the left 55b, the left SCEF, bilateral LO2, and right PF were identified as cross-domain regions. According to previous literature, the left SCEF contributes to the control of both oculomotor sequences and action sequences, including the selection, ordering, and monitoring of sequential behaviors (Abzug & Sommer, 2018; Cona & Semenza, 2017; Nachev, Kennard, & Husain, 2008). The SCEF has also been identified as part of the core multiple-demand (MD) network (Assem, Shashidhara, Glasser, & Duncan, 2022; Camilleri et al., 2018), suggesting a role in cognitive processes shared across multiple task domains. Consistent with this, recent fMRI studies show that core MD regions such as the SCEF and adjacent sensory-biased regions including area 55b form interleaved functional zones rather than isolated modules (Assem et al., 2022). Although LO2 is classically defined as part of the visual cortex, converging evidence indicates that it supports higher-level visual representations such as shape, configuration, and object structure (Cichy, Chen, & Haynes, 2011; MacEvoy & Epstein, 2011). Recent encoding model studies further indicate that such high-level visual representations are aligned with representations in large language models and can generalize across different stimulus modalities (Doerig et al., 2025; Tang et al., 2023). However, the LO2 did not show a significant difference between structured test stimuli and unstructured control stimuli (**Figure 2**), suggesting that its contribution may not be specific to structured symbolic processing. Instead, LO2 may reflect higher-level visual or stimulus-format representations that are shared across the language and math tasks. In contrast, the right PF should be interpreted with caution. Although this region survived the conjunction analysis and therefore met our operational definition of a cross-domain region, its representational and connectivity profiles differed from those of the left 55b, SCEF, and LO2. Specifically, the right PF showed weight distributions and connectivity patterns distinct from those of the other cross-domain regions. This suggests that the right PF may not represent a core region shared between language and mathematics. Instead, it may reflect a parietal interface supporting symbolic manipulation, quantity-related processing, or attentional control that is recruited by both domains to different degrees.

The language specificity and math specificity analyses revealed involvement of the temporal and parietal areas, respectively. These results are consistent with previous studies reporting that language and mathematics are supported by distinct neural networks (Fedorenko et al., 2024; Fedorenko & Varley, 2016; Mahowald et al., 2024): language primarily involves fronto-temporal areas, whereas mathematics engages the fronto-parietal areas. Within the frontal cortex, the middle frontal gyrus has been associated more strongly with mathematics (Fedorenko et al., 2024; Friederici et al., 2017; Price, 2012), while the inferior frontal gyrus has been linked with language (Arsalidou et al., 2018; Nieder, 2025; Nieder & Dehaene, 2009). It should be noted, however, that these findings are not contradictory to our encoding model results suggesting partially shared neural representations for language and mathematics. The encoding model likely captures both shared and domain-specific information, and our analyses captured distinct contributions from these components. Indeed, the weight-correlation analysis and PCA suggested that the latent feature space contained dimensions preferentially associated with cross-domain and domain-specific prediction patterns. Among the top three PCs, PC1 was most strongly associated with the conjunction map, PC2 with the math-specificity map, and PC3 with both the language-specificity and conjunction maps. This property of the latent LLM features may have enabled the construction of an encoding model that simultaneously incorporates both the shared and domain-specific aspects of language and mathematics.

The functional connectivity analyses revealed that, even from the same shared ROIs, connectivity with different networks varied depending on the task domains. The temporal areas that showed stronger connectivity during the language task corresponded to regions with higher language specificity in prediction accuracy. Similarly, the parietal areas that exhibited stronger connectivity during the mathematics task corresponded to regions with higher math specificity in prediction accuracy. These findings suggest that the left 55b, left SCEF, right PF, and LO2 exhibit task-dependent connectivity with language– and mathematics-related networks. Indeed, a diffusion MRI study has reported that the left 55b is anatomically connected to the AIP and pSTS via the second division of the superior longitudinal fasciculus (SLF II) (Poologaindran, Lowe, & Sughrue, 2021), suggesting consistency between anatomical and functional connections among these regions. In addition, the functional connectivity analysis with predicted responses suggested that these connectivity patterns were partly captured by the latent LLM features rather than solely reflecting general cognitive factors unrelated to language or mathematics. Although the present analyses cannot determine whether these task-dependent correlation patterns reflect causal interactions between regions, the seed-by-task interaction analysis showed that the same pattern was evident when seed-target associations were modeled as a function of task domain. Moreover, the predicted-response connectivity analysis suggests that these patterns were partly supported by representational components captured by the latent LLM features.

Several limitations should be noted regarding the present approach. First, the structural variety of sentences and mathematical expressions was limited. We included only mathematical expressions containing two operations, and no stimuli with a single operation (e.g., “2 + 3”). This design was based on the observation of limited prediction performance in the single-operator problems in our previous study (Nakai & Nishimoto, 2023a). Therefore, it remains unclear whether the left 55b and other regions also play roles when participants perform single-operator problems. It is also unclear how activations in this region, if they reflect processing of structured symbol sequences, are modulated by the complexity of structural relationships. Previous studies have reported increased brain activity in the left IFG, a region located in close proximity to the left 55b, as a function of the syntactic complexity of sentences and mathematical expressions (Nakai & Sakai, 2014; Pallier, Devauchelle, & Dehaene, 2011). Although we were unable to include a sufficient number of stimuli to fully assess the structural diversity of language and mathematics in the present study, it will be necessary for future research to conduct a more detailed examination of the structural organization of language and mathematical expressions (Matsumoto & Nakai, 2023).

Second, it would be worth considering the influence of other cognitive factors that may contribute to the shared representations observed between language and mathematics. Regarding simple visual processing, the fact that brain activity could not be sufficiently predicted using AlexNet features suggests that the prediction in the cross-domain regions does not reflect low-level visual information. As for motor components related to button responses, the data from the 20% of trials with button presses were excluded from the analysis, thereby eliminating their influence. Simple short-term memory shared between words and symbols is unlikely to explain the activity in these regions, as its contribution would also appear in the control condition in which word/symbol lists were presented. With respect to more complex working memory processes, some studies have proposed that working memory itself contributes to processing structured symbolic information (Dehaene et al., 2022; Desbordes, King, & Dehaene, 2024; Fitch & Martins, 2014). A detailed examination of these cognitive factors will be an important direction for future research.

In conclusion, the present study provides evidence for partially shared and domain-specific neural representations across language and mathematics. We identified cortical regions showing cross-domain prediction and task-dependent connectivity with language– and mathematics-related networks. Together, these findings help reconcile contrasting views on the neural organization of language and mathematics.

## Methods

### Subjects

Thirty-two subjects, aged 19–30 years (13 females, all with normal vision), denoted as ID01–ID32, participated in this study. Although the present study used subject-wise encoding models, statistical inference was performed at the group level by testing prediction accuracy across participants. We therefore set the target sample size to at least 30 participants, taking into account typical sample sizes used in the neuroimaging literature (Szucs & Ioannidis, 2020). All participants were native Japanese speakers and right-handed (laterality quotient, 30–100) (Oldfield, 1971). Written informed consent was obtained from all subjects before their participation, and the study received approval from the ethics and safety committee of the National Institute of Information and Communications Technology in Osaka, Japan.

### Stimuli and procedures

The current experiment comprised four conditions: a language target condition, a language control condition, a math target condition, and a math control condition. Stimuli in the language target condition consisted of 200 sentences, 160 of which were used for model training (referred to as *language training stimuli*) and 40 for model testing (referred to as *language test stimuli*). Each sentence contained five Japanese words and expressed a complex noun phrase with a relative clause (e.g., *“risu-wo otta neko-ga hei-kara ochita”* [squirrel-ACC chase-PAST cat-NOM fence-ABL fall-PAST]: “The cat that chased the squirrel fell off the fence”). The sentences followed a consistent syntactic structure, with variations in case particles (e.g., *wo*, *de*, *ni*, *kara*) depending on the semantic roles required by the verbs.

Stimuli in the language control condition consisted of 40 word lists, all of which were used only for model testing (referred to as *language control stimuli*). Each word list comprised five unrelated words, consisting either only of nouns or only of verbs, that did not form a grammatical sentence. In the noun lists, each noun was accompanied by a case particle, whereas the verb lists consisted of inflected verbs (e.g., “*kiita, oeta, naoshita, kaita, ushinatta*” [hear.PAST finish.PAST cure.PAST write.PAST lose.PAST]). Thus, the language control condition was designed to be comparable to the language target condition at the word level, while minimizing sentence-level grammatical structure.

Stimuli in the math target condition consisted of 200 mathematical expressions, 160 of which were used for model training (referred to as *math training stimuli*) and 40 for model testing (referred to as *math test stimuli*). Each expression contained five symbols (three single-digit numbers and two operators) and represented a two-operator arithmetic problem (e.g., “7 × 6 + 3”). The expressions followed a consistent structure, with variations in operators (“+”, “−”, “×”, and “/”).

Stimuli in the math control condition consisted of 40 digit lists, all of which were used only for model testing (referred to as *math control stimuli*). Each list contained five isolated single digits that did not form a calculable mathematical expression (e.g., “1, 8, 2, 7, 5”). Operators were not included in this condition because the number of operator types was limited.

At the beginning of each run, a small rectangular cue was presented for 500 ms at the center of the screen, followed by a target sentence or mathematical expression consisting of five words or symbols. Each word or symbol was presented sequentially at the center of the screen for 700 ms, resulting in a total presentation time of 3,500 ms.

In 20% of the trials, the sentence or mathematical expression was followed by a blank screen for 1,000–2,000 ms and then by a probe stimulus presented for 2,000 ms. In the language target condition, participants were instructed to indicate whether the probe matched the meaning of the preceding sentence by pressing the left or right button with their right hand. For example, after the sentence “*risu-wo otta neko-ga hei-kara ochita*” [The cat that chased the squirrel fell off the fence], the probe “*neko-ga ochita*” [The cat fell] was judged as matching the sentence meaning. In the language control condition, participants were instructed to indicate whether the probe word had appeared in the preceding word list. For example, after the word list “*kiita, oeta, naoshita, kaita, ushinatta*” [hear.PAST finish.PAST cure.PAST write.PAST lose.PAST], participants judged whether the probe “kaita?” [write.PAST?] had been included in the list. In the math target condition, participants were instructed to indicate whether the probe digit matched the result of the preceding mathematical expression (e.g., “=21?”) by pressing the left or right button with their right hand. In the math control condition, participants instead indicated whether the probe digit had appeared in the preceding digit list (e.g., “1, 8, 2, 7, 5” followed by “2?”). Task engagement was assessed based on accuracy for trials containing a probe stimulus (see **Supplementary Information**). Following the response, a feedback stimulus was displayed for 1,000 ms, indicating whether the participant’s answer was correct or incorrect. The next run began after a 1,000–2,000 ms intertrial interval. Thus, the duration of a single trial without a probe was 5,000–6,000 ms, whereas trials with a probe lasted 9,000–11,000 ms.

Each run consisted of 40 trials (regardless of condition). At the beginning of each run, 6 s of dummy scans were acquired during which a fixation cross was displayed; these dummy scans were later excluded from the final analyses to reduce noise. An additional 6 s of scans were acquired at the end of each run, during which the fixation cross was also displayed; these scans were included in the analyses. The total duration of each run was 268 s. Each participant completed twelve fMRI runs in total. Of these, five runs were assigned to the language target condition, including four language training runs and one language test run. Another five runs were assigned to the math target condition, including four math training runs and one math test run. The remaining two control runs were assigned to the language and math control conditions, using the corresponding control stimuli. The language and math runs were interleaved, and the run order was reversed for half of the participants.

Stimuli were presented on a projector screen inside the scanner (21.0 × 15.8° of visual angle at 30 Hz). Subjects were equipped with MR-compatible ear tips. Presentation software (Neurobehavioral Systems, Albany, CA, USA) was used to control the stimulus presentation and the collection of behavioral data. To measure button responses, an optic response pad with two buttons held in the right hand was used (HHSC-2 × 2, Current Designs, Philadelphia, PA, USA).

### MRI data acquisition

The experiment was conducted using a 3.0 T scanner (MAGNETOM Vida; Siemens, Erlangen, Germany) with a 64-channel head coil. We scanned 72 interleaved axial slices, 2-mm thick without a gap, parallel to the anterior and posterior commissure line, using a T2*-weighted gradient-echo multiband echo-planar imaging sequence [repetition time (TR) = 1,000 ms; echo time (TE) = 30 ms; flip angle (FA) = 62°; field of view (FOV) = 192 × 192 mm^2^; resolution = 2 × 2 mm^2^; multiband factor = 6]. We obtained 268 volumes in each run. As an anatomical reference, high-resolution T1-weighted images of the whole brain were acquired from all subjects with a magnetization-prepared rapid acquisition gradient-echo sequence (TR = 2,530 ms; TE = 3.26 ms; FA = 9°; FOV = 256 × 256 mm^2^; voxel size = 1 × 1 × 1 mm^3^).

### Preprocessing

The preprocessing of both functional and anatomical MRI data was carried out using fMRIPrep 21.0.2 (Esteban et al., 2019). A B0 fieldmap was derived from phase-drift maps, which were measured using two successive gradient-recalled echo acquisitions. The corresponding phase maps were unwrapped using prelude (FSL 6.0.5.1).

The T1-weighted image underwent a correction for intensity non-uniformity, followed by skull-stripping and tissue segmentation. Brain surfaces were reconstructed using recon-all (FreeSurfer 6.0.1) (Dale, Fischl, & Sereno, 1999). Volume-based spatial normalization to the standard space (MNI152NLin6Asym) was achieved through nonlinear registration with antsRegistration (ANTs 2.3.3).

For functional scans, the first six volumes acquired at the beginning of each run were discarded before preprocessing to allow the MR signal to reach steady state. A reference volume and its skull-stripped version were initially produced using the custom methodology of fMRIPrep. Prior to spatiotemporal filtering, head-motion parameters, including six rotation and translation parameters, were estimated in relation to the reference scan using mcflirt (FSL 6.0.5.1) (Jenkinson, Bannister, Brady, & Smith, 2002). All functional scans underwent slice-time correction to the middle slice, and the reference scan was co-registered with the T1w reference using bbregister (FreeSurfer) (Greve & Fischl, 2009). Three global signals were extracted within the cerebrospinal fluid (CSF), white matter (WM), and whole-brain masks. The blood oxygenation level-dependent time-series was resampled into a standard space (MNI152NLin6Asym). All resampling was conducted using a single interpolation step by composing all the relevant transformations (i.e., head-motion transform matrices, susceptibility distortion correction, and co-registrations to anatomical and output spaces).

The resulting functional data underwent high-pass filtering at 0.01 Hz and low-pass filtering at 0.15 Hz. Confounding factors, including six head-motion parameters, global signals from the CSF, WM, and whole-brain masks, as well as the derivative of each factor, were eliminated using linear regression implemented in Scikit-learn. The functional data were then standardized to have a zero mean and unit variance for each of the 18,715 cortical vertices in the fsaverage5 space, excluding non-cortical vertices. To remove motor components related to button responses, we excluded volumes corresponding to trials with button presses and all volumes within the subsequent 6 s, corresponding to the maximum temporal delay used in the encoding model. After this exclusion procedure, each set of four language or math training runs comprised 823–831 samples, depending on the participant. Each language or math test run and each control run comprised 205–213 samples, depending on the participant.

### Feature extraction

Latent LLM features were extracted using the Japanese version of the Llama3 model (https://huggingface.co/elyza/Llama-3-ELYZA-JP-8B) (Dubey et al., 2024), a transformer-based LLM. For each input stimulus, the hidden states (i.e., block outputs after residual connections) from the 1^st^, 8^th^, 16^th^, 24^th^, and 32^nd^ layers (out of a total of 32 layers) were extracted as latent features. For each target layer, the latent features from the language and math training stimuli were concatenated and then reduced to 100 dimensions using PCA (here referred to as *joint PCA projection*; **Figure 1A**). To avoid any information leakage from the test stimuli, PCA was fitted only on the training stimuli, and the resulting PCA projection was then applied to the independent test stimuli and unstructured control stimuli. As an additional analysis, we prepared 100-dimensional features by separately applying PCA for language and math training stimuli (without concatenation; here referred to as *separate PCA projection*; **Supplementary Figure S2C**). Because these analyses relied on a common projection space for language and math features, we verified that joint PCA projection preserved substantial within-domain variance in both domains. The first 100 joint PCs retained 63.1% of the variance in language features and 99.4% in math features, with only modest reductions using separate PCA projection (**Supplementary Information**).

To check reproducibility, we extracted latent LLM features from other models, including 1^st^, 6^th^, 13^th^, 20^th^, and 26^th^ layers of Gemma2 (https://huggingface.co/google/gemma-2-2b-jpn-it) (Gemma Team et al., 2024), 1^st^, 7^th^, 14^th^, 21^st^, and 28^th^ layers of Qwen2.5 (Yang et al., 2024), 1^st^, 8^th^, 16^th^, 24^th^, and 32^nd^ layers of Japanese version of GPT-NeoX (https://huggingface.co/docs/transformers/model_doc/gpt_neox_japanese) (Black et al., 2022), 1^st^, 8^th^, 16^th^, 24^th^, and 32^nd^ layers of RakutenAI2 (https://huggingface.co/Rakuten/RakutenAI-2.0-8x7B), and 1^st^, 10^th^, 20^th^, 30^th^, and 40^th^ layers of LLM-jp (LLM-jp et al., 2024).

To further examine sublayer-specific contributions within the transformer, we additionally extracted intermediate representations from the 16th layer. Specifically, we obtained the outputs of the self-attention and MLP sublayers prior to residual addition. These sublayer-specific features were analyzed separately to assess their respective contributions to shared and domain-specific representations across input elements.

As a control for low-level visual processing, we extracted latent visual features using AlexNet, a convolutional neural network widely used to model visual processing in the human brain (Gu et al., 2022; A. Y. Wang et al., 2023; Wen et al., 2018). For each visual stimulus, the output from the first convolutional layer was extracted, and the latent features from the language and math training stimuli were concatenated and then reduced to 100 dimensions using PCA.

### Vertex-wise encoding model fitting

In the encoding model, each vertex’s cortical activity was modeled using a finite impulse response model. This model captured the slow hemodynamic response and its association with neural activity (Kay, David, Prenger, Hansen, & Gallant, 2008; Nishimoto et al., 2011). A feature matrix of dimensions [T × 4N] was constructed by concatenating sets of [T × N] feature matrices, with four temporal delays ranging from 3 to 6 s (where T = number of samples; N = number of features). The cortical response of dimensions [T × V] was then modeled by multiplying the feature matrix by the weight matrix of dimensions [4N × V] (where V = number of vertices). The weight matrix was obtained using L2-regularized linear regression with the training samples. The optimal regularization parameter was determined using 5-fold cross-validation, where the regularization parameters varied from 1 to 10^6^ across seven different values. As a result, two encoding models were obtained for each subject. These models were denoted as the *language encoding model* and the *math encoding model* based on their training domains (language and mathematics), respectively.

The prediction accuracy was evaluated using Pearson’s correlation coefficient between the predicted and actual test samples. Statistical significance (one-sided) was calculated for each vertex using a Wilcoxon signed-rank test (*p* < 0.05), and multiple comparisons were corrected using the false discovery rate (FDR) correction method (Benjamini & Hochberg, 1995). To assess the robustness of the prediction results, we also performed a block-permutation test (see **Supplementary Information**). Prediction accuracy was computed in both intra-domain and cross-domain settings. In the intra-domain prediction, the language encoding model was tested with the language test samples, or the math encoding model was tested with the math test samples. In the cross-domain prediction, a language encoding model was tested with the math test samples, or the math encoding model was tested with the language test samples. Furthermore, the conjunction analysis was performed using the minimum prediction accuracy across all prediction types (i.e., language to language, math to math, language to math, and math to language) (Nichols et al., 2005). This analysis is expected to identify brain regions that represent information shared between language and mathematics. For visualizing data on cortical maps, we used pycortex (Gao, Huth, Lescroart, & Gallant, 2015).

Our main analysis computed prediction accuracy for the language and math target conditions using LLM (Llama3) features with joint PCA projection. In addition, we performed three control analyses: (1) prediction accuracy for the language and math control conditions using LLM features with joint PCA projection, (2) prediction accuracy for the language and math target conditions using visual (AlexNet) features with joint PCA projection, and (3) prediction accuracy for the language and math target conditions using LLM features with separate PCA projection.

### Principal component analysis

To show the structure of the representational space shared by language and mathematics, we performed PCA on the weight matrix concatenated across the language and math encoding models. The 100-dimensional latent features were mapped onto the two-dimensional space using the loadings of the second and third PCs, i.e., PC2 and PC3, as the x-axis and y-axis, respectively. In this space, the operators were colored red, green, and blue based on the relative PCA loadings in PC2, PC1, and PC3, respectively. To visualize the cortical organization, we extracted and normalized the PCA scores from each vertex. The resulting cortical map indicated the relative contribution of each cortical vertex to the top three PCs (PC2, red; PC1, green; PC3, blue).

### Functional connectivity analyses

To elucidate how the brain regions associated with mathematics and language, identified through the basic prediction accuracy analysis of the encoding model, are functionally related to language-specific and math-specific regions, we conducted two types of analyses. First, in the functional connectivity analysis, the Pearson correlation coefficient was calculated between the activity time course extracted from a seed anatomical ROI (averaged across vertices within the ROI), and that of each vertex and each of the remaining anatomical ROIs across the cortex for each training run of the math target and the language target conditions. Functional connectivity was then defined as the mean correlation coefficient across training runs; one functional connectivity map was then obtained for each of the language and math conditions.

Second, to test whether the functional connectivity is indeed related to the latent representations captured by encoding models, we performed a similar analysis using predicted brain responses by the intra-domain encoding models. Pearson correlation coefficient was calculated between the predicted responses extracted from a seed anatomical ROI (averaged across vertices within the ROI) and those of each cortical vertex and each of the remaining anatomical ROIs for both the language and math encoding models.

For both analyses, vertices showing significantly larger functional connectivity in the language condition than in the math condition, as well as those showing the opposite pattern, were examined using a Wilcoxon signed-rank test (*p* < 0.05) with the FDR correction. ROI-based functional connectivity results were visualized using the MNE-Connectivity Python package (Gramfort et al., 2014). The 360 ROIs in the original HCP-MMP1 atlas were integrated into 21 left-hemisphere and 21 right-hemisphere regions, following the grouping provided at https://neuroimaging-core-docs.readthedocs.io/en/latest/pages/atlases.html. The “Primary Visual” region was included in the “Early Visual” region.

## Acknowledgements

We thank JSPS KAKENHI (grant numbers 26H00539 for T.N., JP18H05522 and JP24H00619 for S.N.), JST ERATO (JPMJER1801), JST AIP (JPMJCR24U2 for S.N.), JST FOREST Program (JPMJFR231V for T.N.), and JST Moonshot R&D (JPMJMS259G for T.N.) for partial financial support of this study. The funders had no role in the study design, data collection and analysis, decision to publish, or preparation of the manuscript.

## Author contributions

**T.N.**: Conceptualization, Methodology, Formal analysis, Visualization, Writing – original draft preparation. **R.K.**: Data collection, Writing – Review & Editing. **S.N.**: Supervision, Writing – Review & Editing, Funding acquisition.

## Supplementary Information

### Behavioral data analysis

To confirm that participants were sufficiently engaged with the experimental tasks, we analyzed accuracy for trials in which a probe stimulus was presented (20% of all trials). Participants performed both the language and math target conditions with high accuracy (>90%). In the language domain, accuracy was significantly higher in the target condition than in the control condition (Target: M = 0.912, SD = 0.046; Control: M = 0.863, SD = 0.118; Wilcoxon signed-rank test, *p* = 0.039). In contrast, no significant difference was observed between the target and control conditions in the math domain (Target: M = 0.958, SD = 0.049; Control: M = 0.968, SD = 0.064; *p* = 0.252). Thus, the control conditions were not easier than the corresponding target conditions; if anything, the language control condition was more difficult than the language target condition.

### Block-permutation test

To further assess whether the cross-domain prediction results were statistically reliable while accounting for the temporal structure of the fMRI responses, we performed a block-permutation test. For each participant and prediction type, the test-run time series was divided into consecutive blocks of 10 samples. The order of these blocks was then randomly permuted, thereby disrupting the temporal correspondence between the predicted and observed responses while preserving the local temporal structure within each block. Prediction accuracy was recalculated after each block permutation.

This procedure was repeated 5,000 times. For each permutation, prediction accuracies were averaged across participants to generate a null distribution of group-mean prediction accuracy at each cortical vertex. The observed group-mean prediction accuracy was then compared with this permutation-based null distribution. A one-sided *p* value was computed as the proportion of permutations in which the group-mean prediction accuracy was equal to or greater than the observed value. The resulting significance maps were FDR-corrected for multiple comparisons across cortical vertices using the same procedure as in the main analysis, and the conjunction analysis across intra-domain and cross-domain predictions was repeated using these block-permutation-based significance maps.

### Validation of joint PCA as a common feature space

A potential concern with applying joint PCA to language and math features is that a common projection space may fail to preserve the dominant variance structure of one or both domains. To assess whether joint PCA provided an appropriate common feature space for language and math representations, we examined how much variance in each domain was retained after projection onto the joint PCA components. Because joint PCA is fit to the concatenated language and math features, it does not necessarily maximize variance within either domain alone. In contrast, separate PCA provides an upper-bound estimate of how much variance can be captured when dimensionality reduction is optimized independently for each domain. We therefore compared the variance explained by the first 100 components of the joint PCA with that explained by the first 100 components of separate PCAs fit to the language and math features independently.

The first 100 components of the joint PCA explained 63.1% of the variance in language features and 99.4% in math features. In contrast, the first 100 components of separate PCAs explained 70.1% and 99.8%, respectively. As expected, separate PCA preserved more within-domain variance than joint PCA, but the reduction associated with using a common projection space was modest. These results indicate that joint PCA retained substantial variance in both domains while allowing language and math features to be represented in a shared feature space. The particularly high variance explained for math features likely reflects the restricted structure of the math stimuli. In the present experiment, the math stimuli consisted of relatively simple expressions, such as “7 × 6 + 3,” composed of single-digit numbers and two arithmetic operators. Thus, compared with the language stimuli, the math stimuli contained a more limited set of constituent elements, which may have resulted in a lower-dimensional feature structure and consequently higher variance explained by the first 100 components.

### Circular-shift control

To test whether prediction depended on the correct correspondence between stimuli and LLM-derived features, we performed a circular-shift control. For each participant and training run, the feature time series was circularly shifted by a random offset before temporal-delay expansion and before excluding time points associated with motor responses. Because each run contained 262 TRs after downsampling, random offsets were sampled from 30–232 TRs, avoiding shifts close to 0 or the full run length. The same time points excluded in the main analysis were then removed from both the shifted feature matrix and the brain-response matrix. This procedure disrupted the correspondence between stimuli and LLM-derived features while preserving the marginal distribution and run-wise temporal structure of the feature matrix. Encoding models were trained on the shifted training features and evaluated on the original independent test features using the same procedure as in the main analysis. This control analysis was repeated 100 times with different random shifts.

### Residual-response encoding analysis

To examine whether the encoding results were driven by mean task-evoked response components, we repeated the encoding analysis using residualized brain responses. For each run, cortical time series were residualized by fitting task-period boxcar regressors convolved with a canonical hemodynamic response function. The task-period regressor modeled the cue and stimulus presentation period of each trial. Probe and feedback periods were modeled as separate nuisance regressors. Run intercepts, linear trends, and quadratic trends were also included as nuisance regressors. The residual time series were then used as response variables in the same LLM-based ridge regression pipeline as in the main analysis. Time points associated with motor responses were excluded after residualization using the same procedure as in the main analysis. The same residualization procedure was applied to the original test runs, control runs, Visual model analysis, and separate-PCA analysis.

### PCA stability analysis

To assess the robustness of the PCA-based model-weight visualization, we repeated the PCA analysis using a leave-one-subject-out procedure. In each iteration, one participant was excluded, group-averaged model weights were recomputed, and PCA was performed using the same procedure as in the main analysis. The resulting PC maps were compared with the full-sample PC maps using spatial correlations across cortical vertices. Because the sign of principal components is arbitrary, PC signs were aligned to the full-sample PCs before computing spatial correlations.

### Seed-by-task interaction analysis

To further examine whether seed-target associations were modulated by task domain, we performed a seed-by-task interaction analysis. For each participant and each anatomically defined seed ROI, the mean seed time series was extracted from the preprocessed BOLD responses. Time points associated with motor responses were excluded using the same procedure as in the functional connectivity analysis. The remaining time series from the language and math test runs were z-scored within each task and concatenated. Task domain was coded as +1 for the language task and −1 for the math task. For each cortical vertex, we fitted a general linear model including the seed time series, the task-domain regressor, and their interaction term. The interaction coefficient was used as an index of task-dependent modulation of the seed-target association. Positive interaction coefficients indicate stronger seed-target associations during the language task than during the math task, whereas negative coefficients indicate stronger associations during the math task. Group-level significance was assessed across participants using a two-sided Wilcoxon signed-rank test, followed by false discovery rate correction across ROIs.

**Figure S1.**
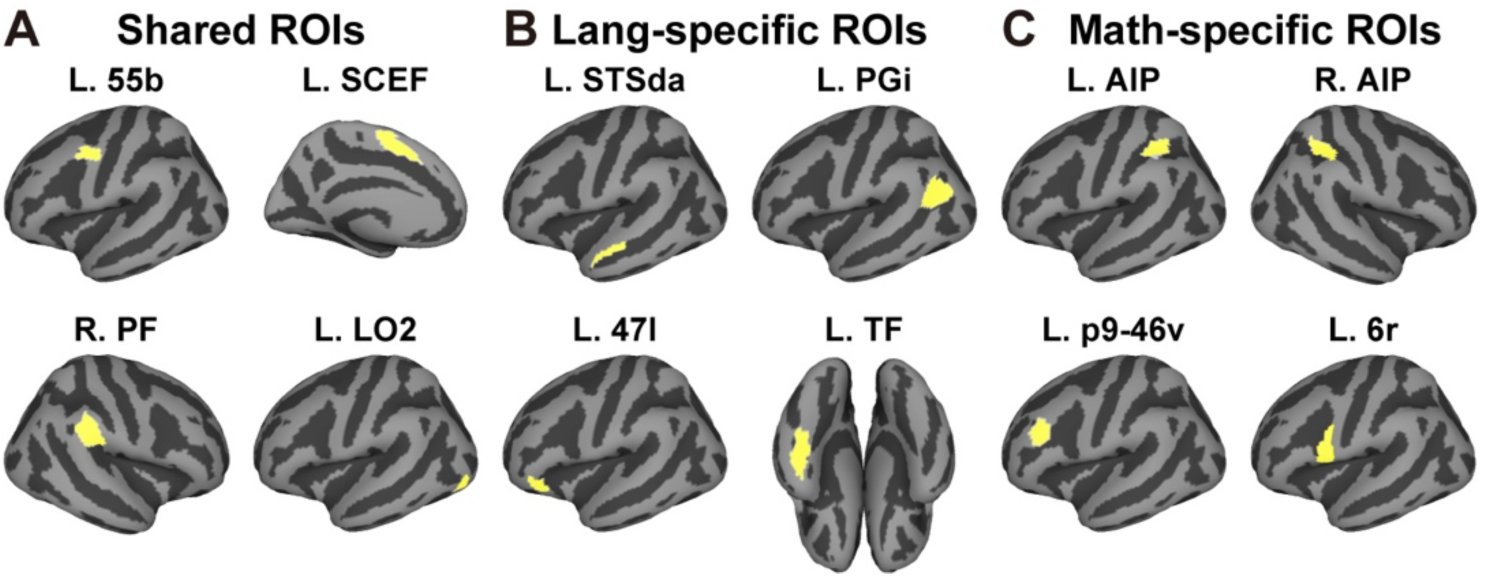
Anatomical regions of interest (ROIs). Target anatomical ROIs of the left 55b, left supplementary cingulate eye field (SCEF), right parietal area F (PF), left area lateral occipital 2 (LO2), left superior temporal sulcus dorsal anterior (STSda), left parietal area G inferior (PGi), left inferior frontal gyrus (47l), temporal fusiform (TF), bilateral anterior intraparietal (AIP), left posterior lateral prefrontal cortex (p9-46v), and left rostral area 6 (6r) are shown in yellow. L, left; R, right.

**Figure S2.**
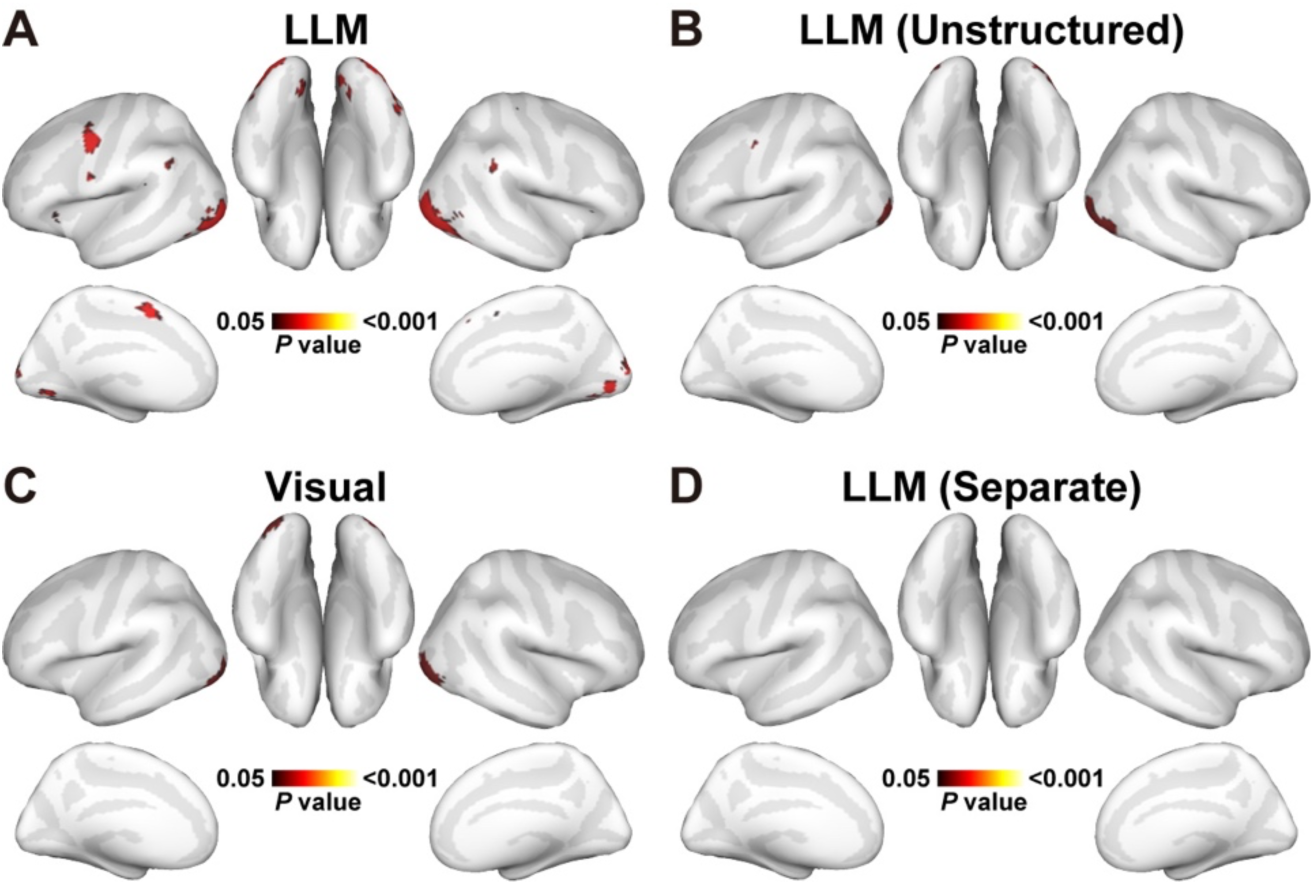
Cross-domain conjunction analysis using block-permutation test. Brain regions significant in both intra– and cross-domain predictions, revealed by conjunction analysis using latent features of **(A)** the original LLM (Llama3) model, **(B)** LLM with unstructured stimuli, **(C)** Visual model, and **(D)** LLM with separate PCA projection.

**Figure S3.**
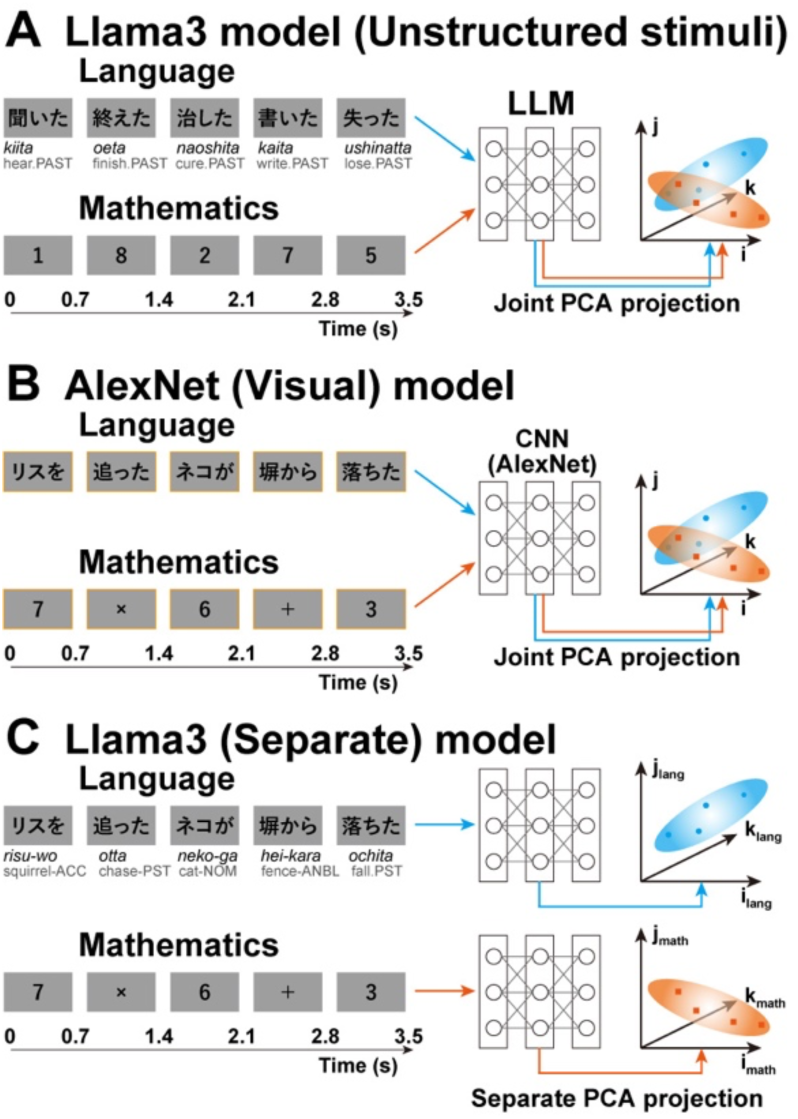
Control model designs. **(A)** Latent features were extracted from both language and math unstructured stimuli using Llama3, embedded into a shared feature space with a principal component analysis (PCA)-based dimensionality reduction. The unstructured stimuli consisted of word or digit lists presented without combinatorial integration. **(B)** Latent features were extracted from both language and math test stimuli using AlexNet, embedding them into a shared feature space. **(C)** Latent features were extracted from both language and math test stimuli using Llama3, embedded into separate feature spaces.

**Figure S4.**
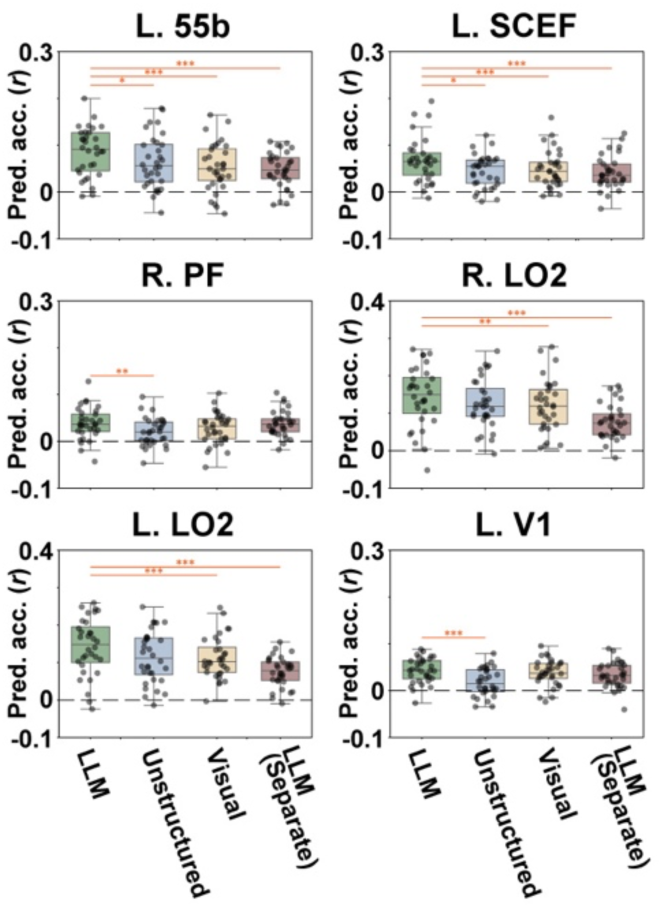
ROI-based prediction accuracy using original brain responses. Box plots show mean prediction accuracy across intra– and cross-domain predictions using original brain responses, without removing task-evoked boxcar components. Prediction accuracies were extracted from anatomical ROIs corresponding to the left 55b, left SCEF, right PF, bilateral LO2, and left V1. Results are shown for the original LLM (Llama3) model, LLM with unstructured stimuli, Visual model, and LLM with separate PCA projection. Dots represent individual participants. Asterisks indicate significance levels in ROI-based comparisons: *, *p* < 0.05; **, *p* < 0.01; ***, *p* < 0.001.

**Figure S5.**
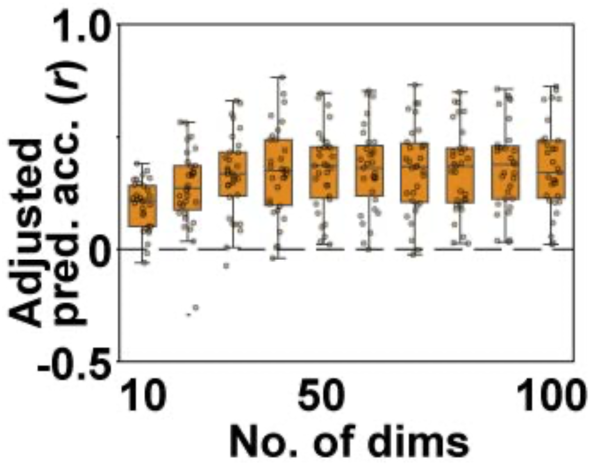
Prediction accuracy with different numbers of dimensions. Box plots present mean prediction accuracy across intra– and cross-domain predictions for Llama3 model, extracted from the left 55b ROI, with an increasing number of dimensions from 10 to 100.

**Figure S6.**
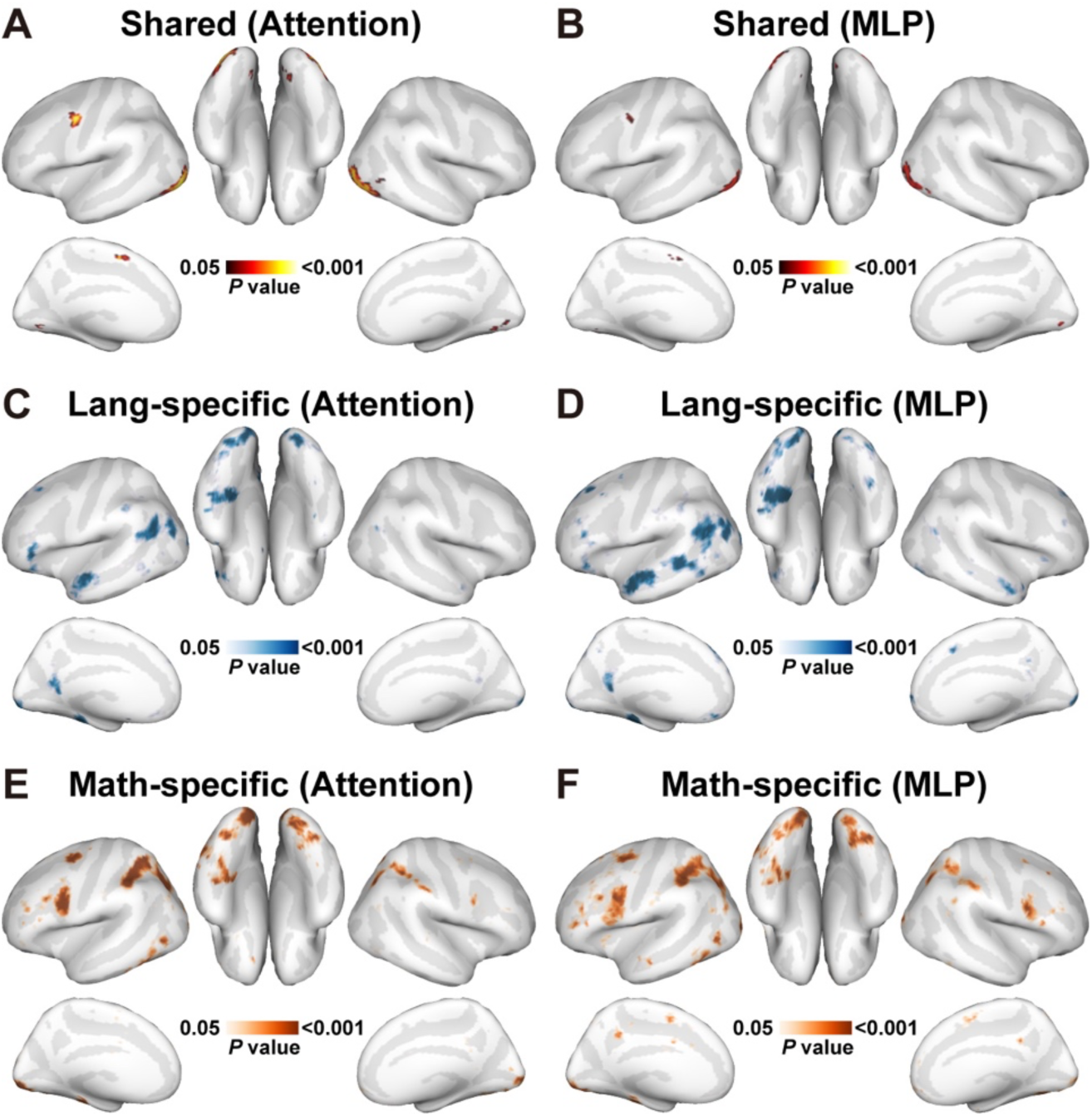
Cross-domain conjunction and domain specificity analyses using attention and MLP sublayers. (**A-B**) Cortical maps showing brain regions significant in both intra– and cross-domain predictions using latent features of **(A)** attention and **(B)** MLP subcomponents of the Llama3 model, thresholded at FDR-corrected *p* < 0.05. **(C-F)** Cortical maps showing brain regions with significantly higher intra-domain prediction than cross-domain prediction for the language target condition using latent features of **(C)** attention and **(D)** MLP sublayers, and for math target condition using latent features of **(E)** attention and **(F)** MLP sublayers.

**Figure S7.**
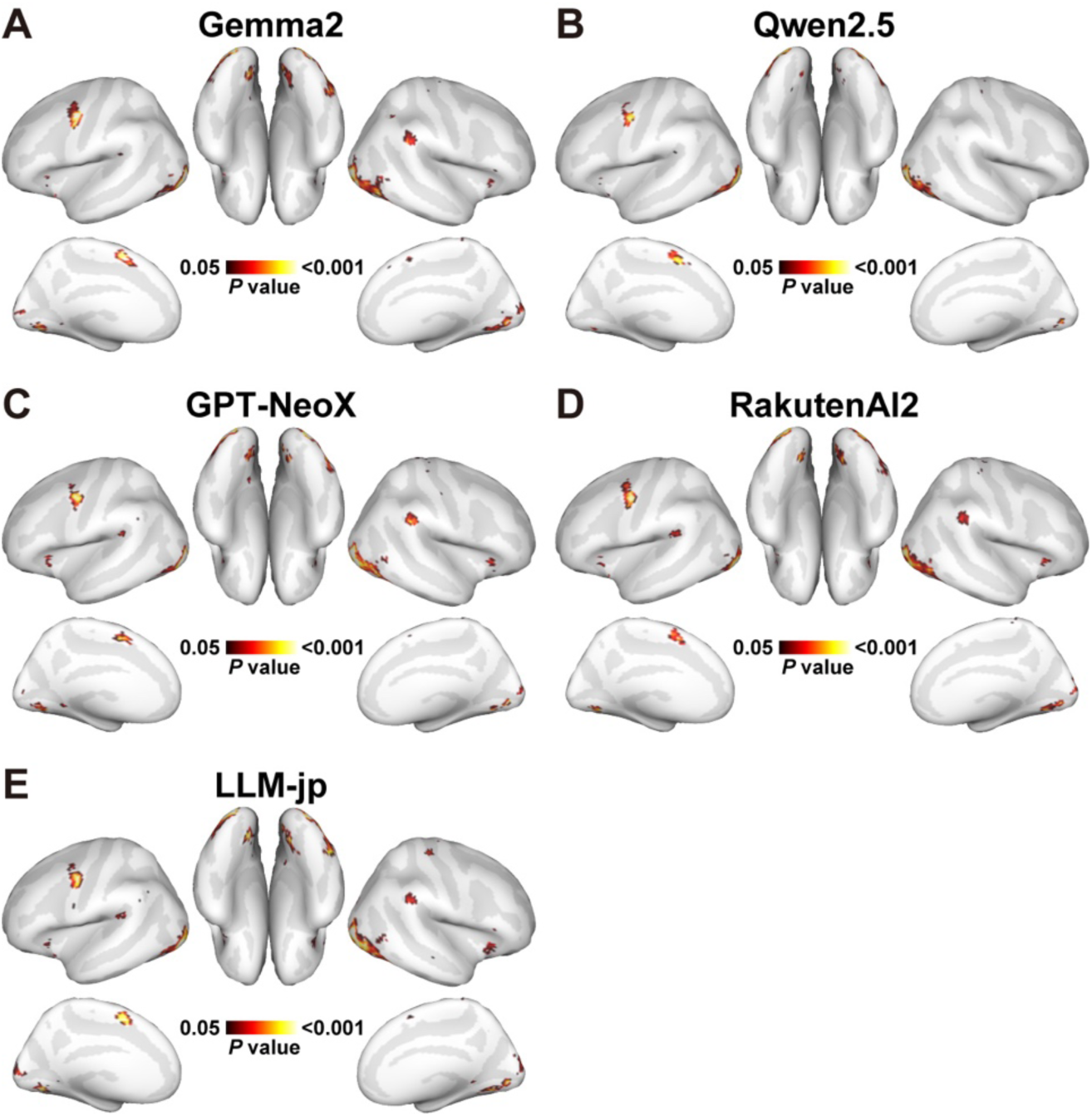
Cross-domain conjunction analysis using other LLMs. Brain regions significant in both intra– and cross-domain predictions, revealed by conjunction analysis using latent features of **(A)** Gemma2, **(B)** Qwen2.5, **(C)** GPT-NeoX, **(D)** RakutenAI2, and **(E)** LLM-jp, thresholded at FDR-corrected *p* < 0.05.

**Figure S8.**
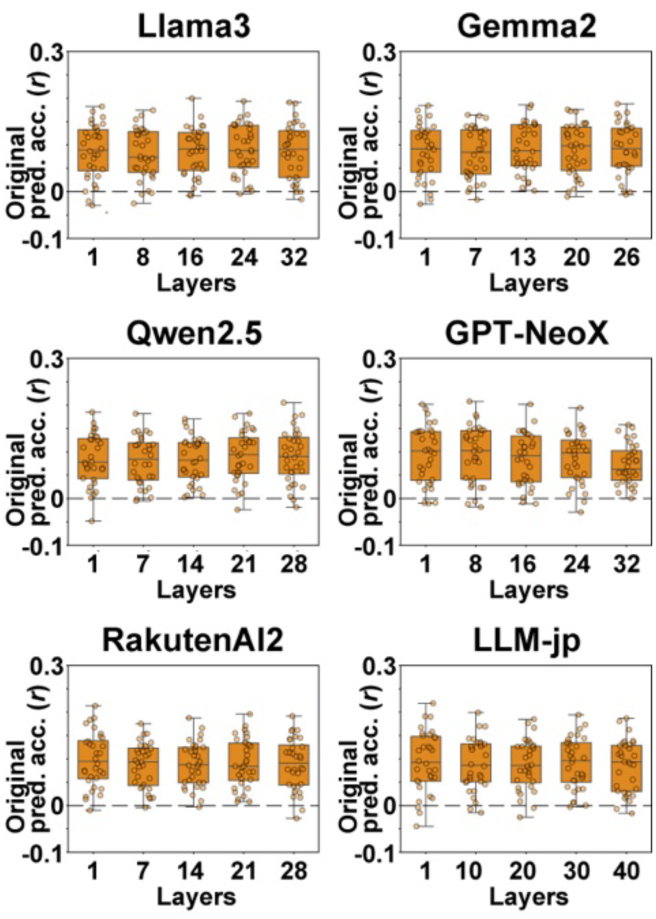
ROI-based cross-domain conjunction analysis using different LLMs. Box plots show mean prediction accuracy across intra– and cross-domain predictions using latent features extracted from five different layers of Llama3, Gemma2, Qwen2.5, GPT-NeoX, RakutenAI2, and LLM-jp.

**Figure S9.**
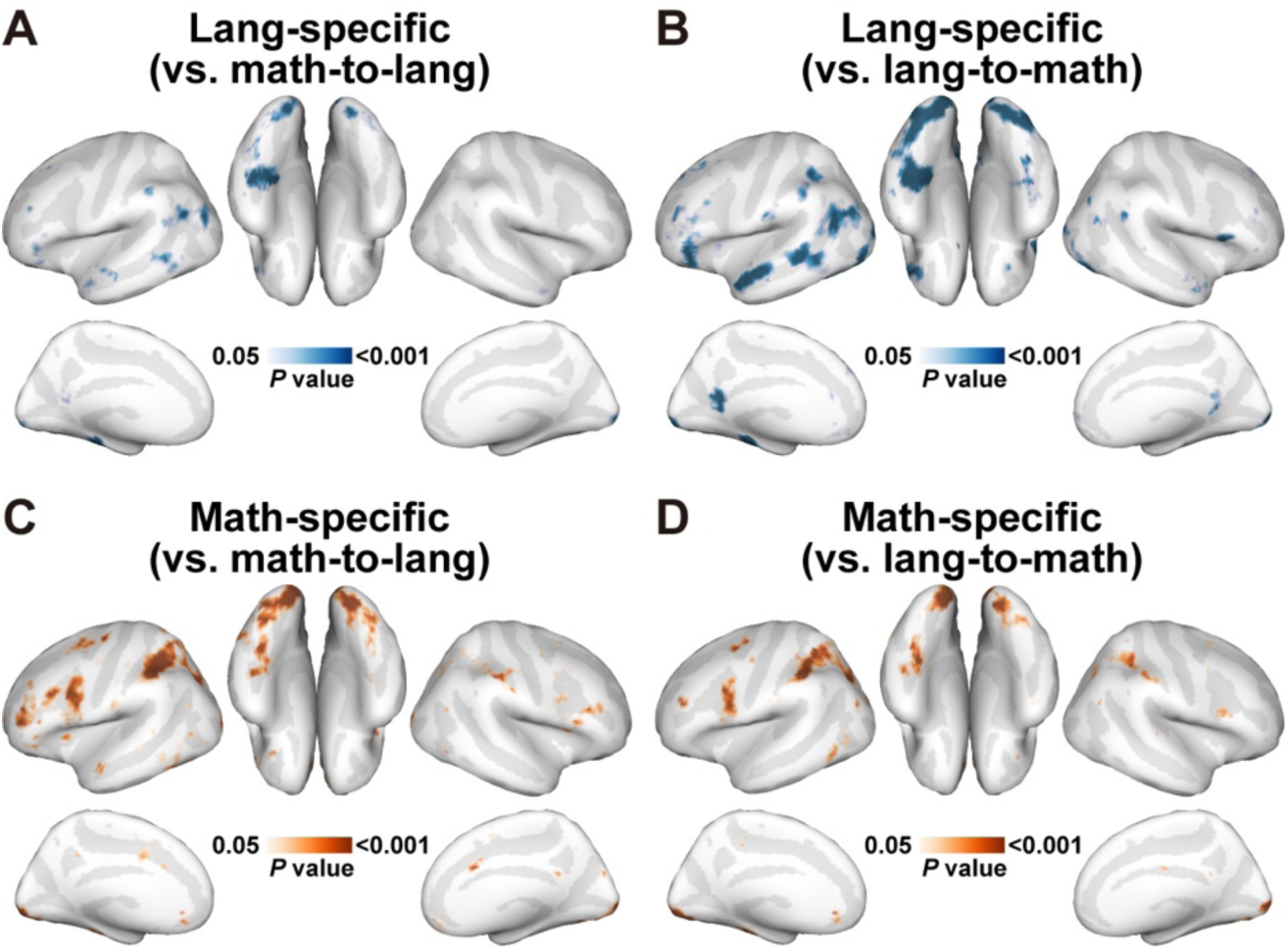
Domain specificity analyses with alternative contrasts. (**A-B**) Language-specificity maps based on contrasts between lang-to-lang and (**A**) math-to-lang or (**B**) lang-to-math prediction, thresholded at FDR-corrected *p* < 0.05. (**C-D**) Math-specificity maps based on contrasts between math-to-math and (**C**) math-to-lang or (**D**) lang-to-math prediction.

**Figure S10.**
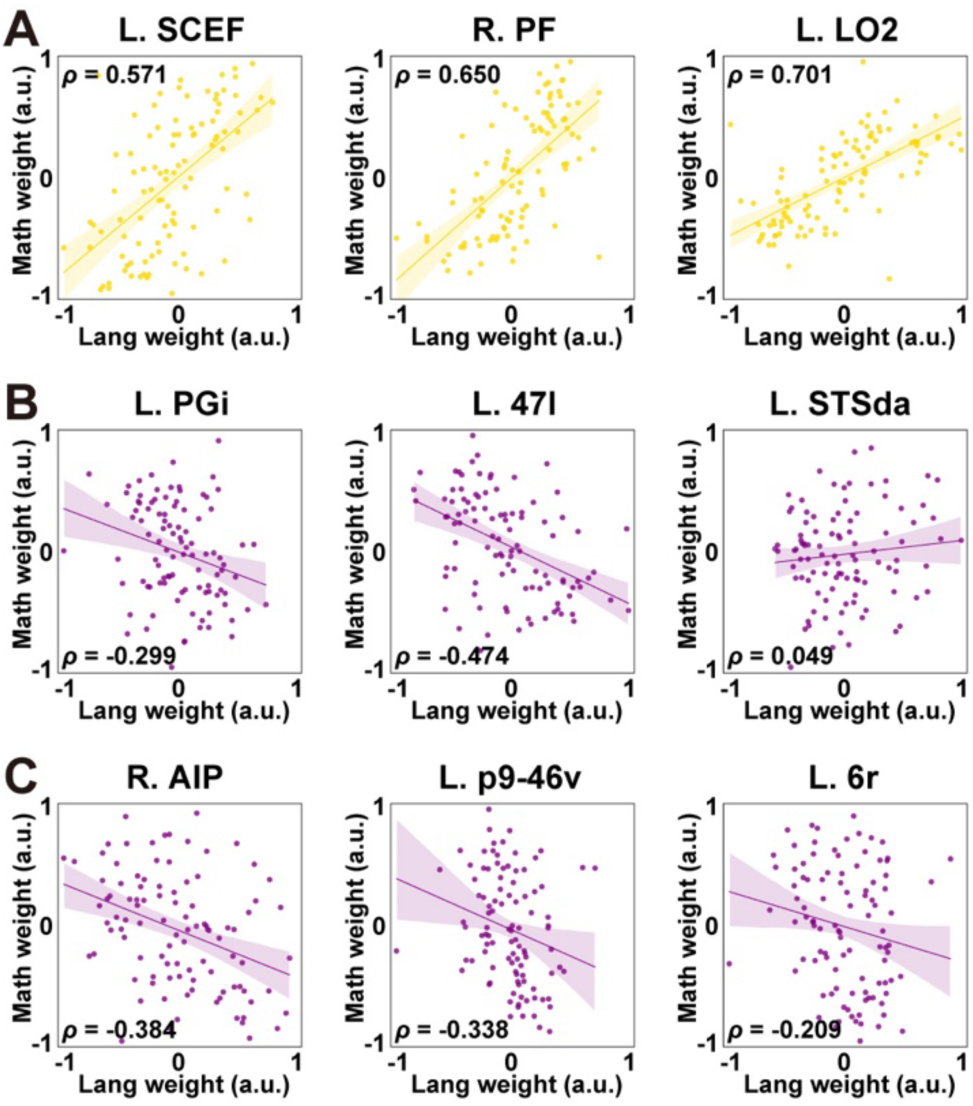
Weight correlation analysis in anatomical ROIs. Scatter plots showing correlation between language and math model weights in the left SCEF, right PF, left LO2, left PGi, left 47l, left STSda, right AIP, left p9-46v, and left 6r.

**Figure S11.**
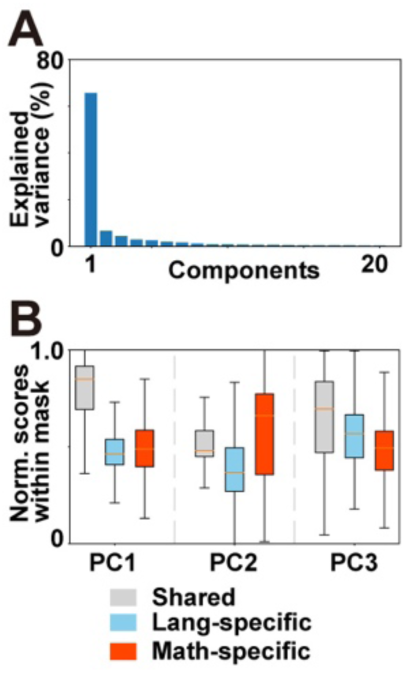
Explained variance and distribution of PC scores. (**A**) Explained variance of top twenty PCs. (**B**) Box plots showing normalized PC scores for the top three PCs, quantified across significant vertices within the conjunction (gray), language-specific (blue), and math-specific (red) maps.

**Figure S12.**
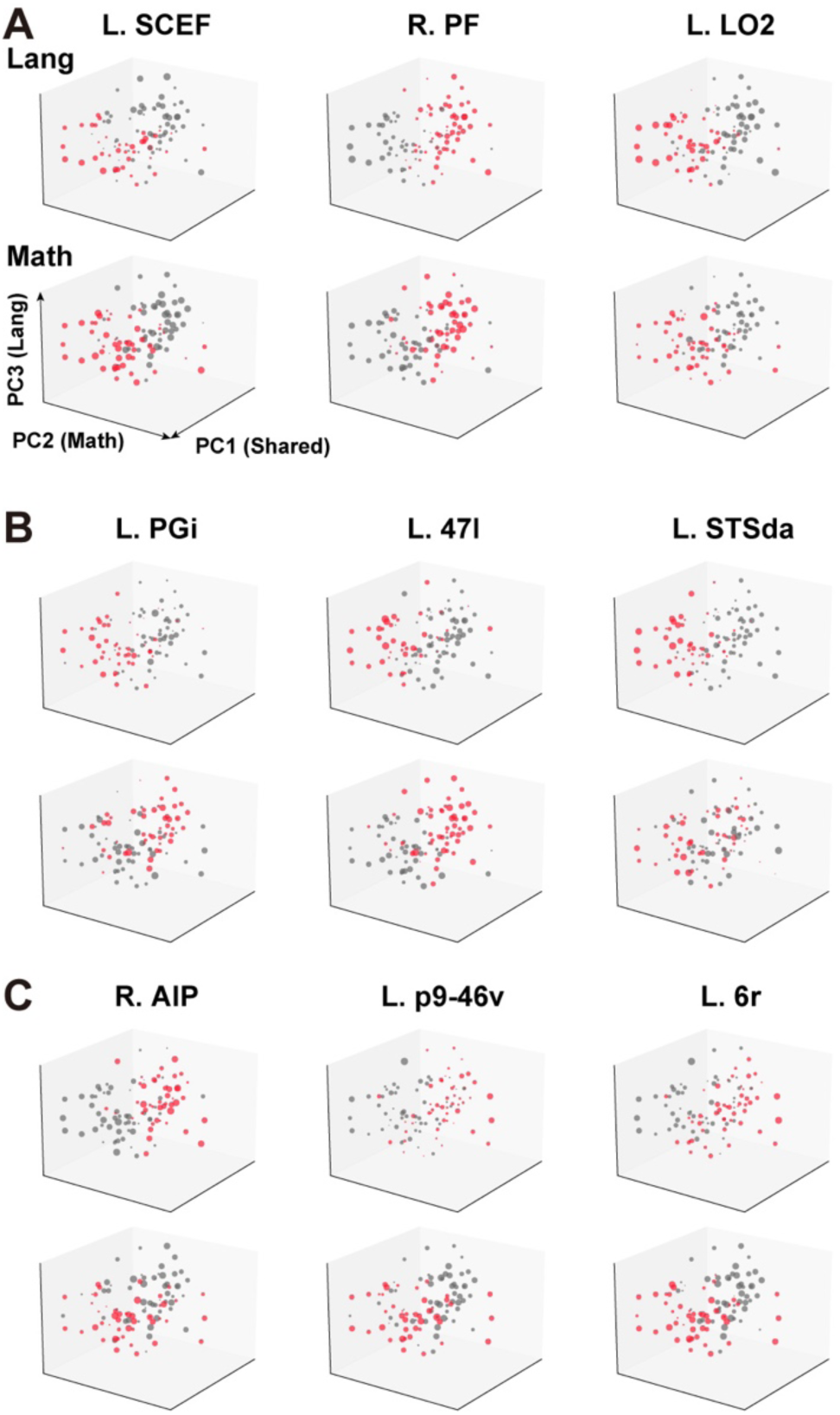
PCA three-dimensional mapping of feature dimensions in anatomical ROIs. Scatter plots showing the weight values of the 100 latent features in the left SCEF, right PF, left LO2, left PGi, left 47l, left STSda, right AIP, left p9-46v, and left 6r, mapped onto a three-dimensional space defined by the PC1, PC2, and PC3 axes.

**Figure S13.**
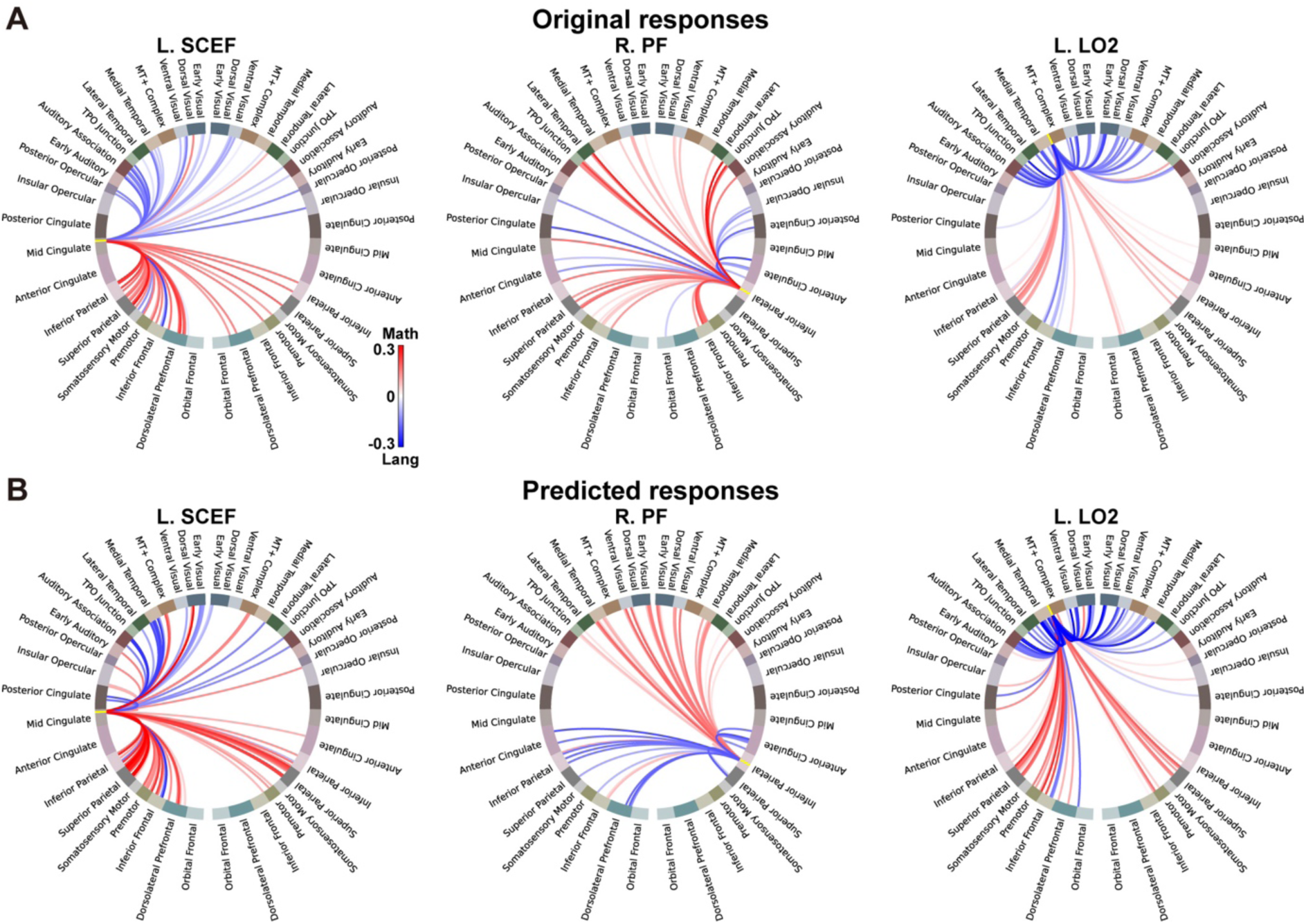
ROI-based model connectivity in the other anatomical ROIs. A circle diagram of functional connectivity computed based on the (**A**) ROI-averaged original responses and (**B**) ROI-averaged predicted responses by the intra-domain encoding models.

**Figure S14.**
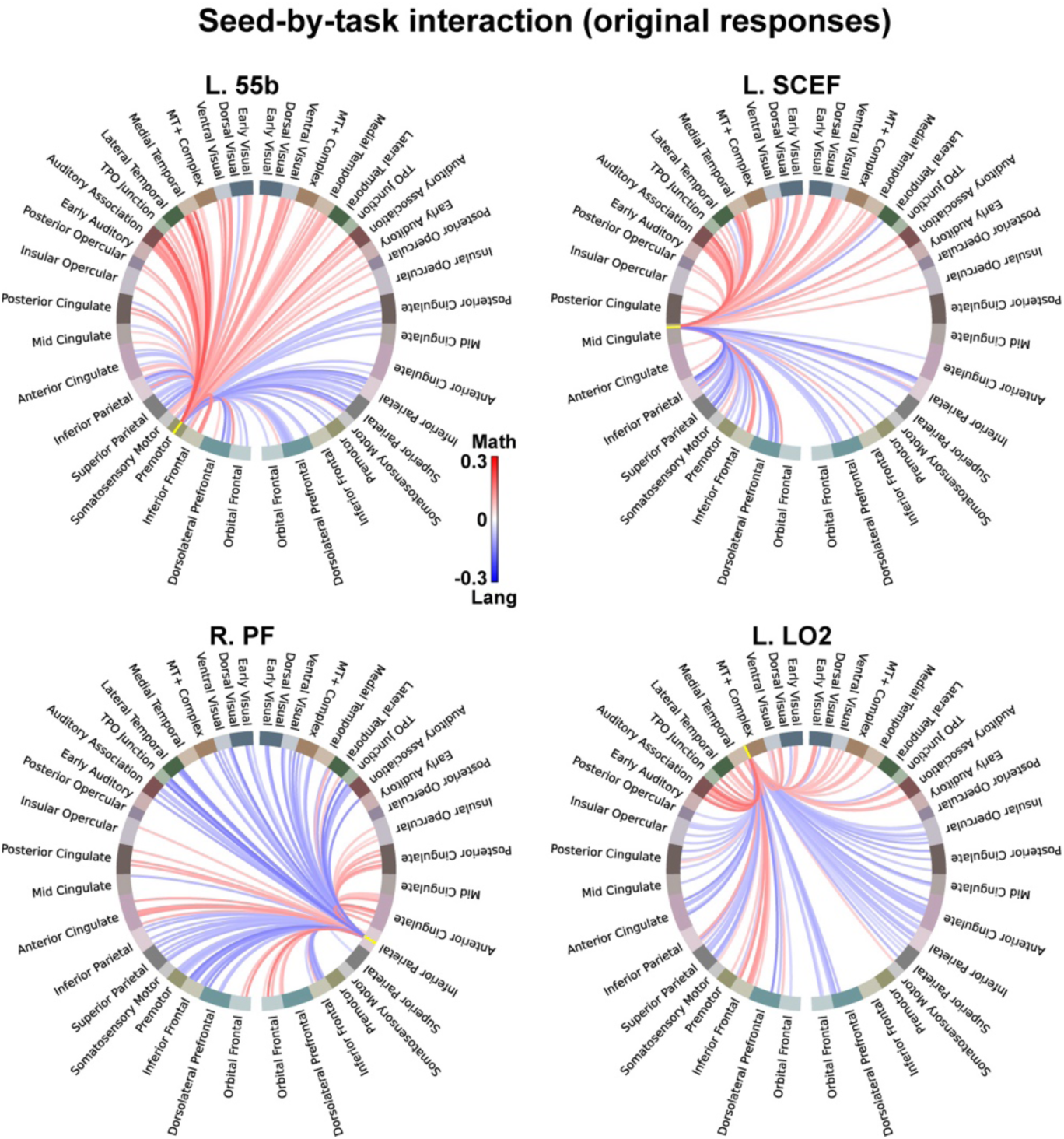
Seed-by-task interaction analysis. Circle diagrams of ROI-based seed-by-task interaction effects using the left 55b, left SCEF, left LO2, and right PF as seed ROIs. ROIs with significantly stronger associations during the language task are shown in red, and those with significantly stronger associations during the math task are shown in blue (two-sided Wilcoxon signed-rank test, FDR-corrected *p* < 0.05). Task-domain main effects were included separately from the seed-by-task interaction term.

**Table S1.**
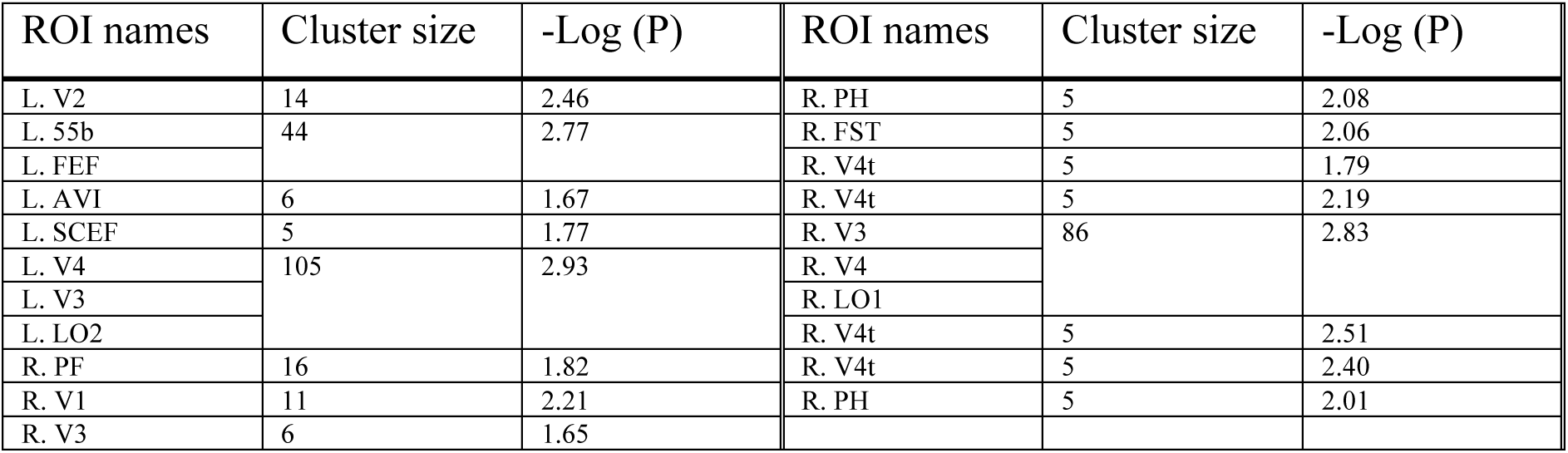
Shared brain regions for language and mathematics. Only clusters with at least 5 vertices are shown.

| ROI names | Cluster size | -Log (P) | ROI names | Cluster size | -Log (P) |
| --- | --- | --- | --- | --- | --- |
| L. V2 | 14 | 2.46 | R. PH | 5 | 2.08 |
| L. 55b | 44 | 2.77 | R. FST | 5 | 2.06 |
| L. FEF |  |  | R. V4t | 5 | 1.79 |
| L. AVI | 6 | 1.67 | R. V4t | 5 | 2.19 |
| L. SCEF | 5 | 1.77 | R. V3 | 86 | 2.83 |
| L. V4 | 105 | 2.93 | R. V4 |  |  |
| L. V3 |  |  | R. LO1 |  |  |
| L. LO2 |  |  | R. V4t | 5 | 2.51 |
| R. PF | 16 | 1.82 | R. V4t | 5 | 2.40 |
| R. V1 | 11 | 2.21 | R. PH | 5 | 2.01 |
| R. V3 | 6 | 1.65 |  |  |  |

**Table S2.** Language-specific brain regions, obtained using Lang-to-Lang vs. average of Lang-to-Math and Math-to-Lang contrast. Only clusters with at least 5 vertices are shown.

| ROI names | Cluster size | -Log (P) | ROI names | Cluster size | -Log (P) |
| --- | --- | --- | --- | --- | --- |
| L. i6-8 | 8 | 2.05 | L. TE2p | 10 | 2.94 |
| L. FOP4 | 10 | 1.91 | L. VVC | 9 | 2.66 |
| L. PFop | 6 | 1.75 | L. PHA2 | 6 | 2.27 |
| L. PFop | 27 | 2.75 | L. PHA3 | 29 | 4.27 |
| L. 7PL | 43 | 2.25 | L. PHA2 |  |  |
| L. POS2 |  |  | L. PHA1 |  |  |
| L. 7Pm |  |  | L. TF | 23 | 4.73 |
| L. v23ab | 8 | 3.37 | L. TF | 117 | 5.06 |
| L. POS1 | 66 | 3.59 | L. PHA3 |  |  |
| L. v23ab |  |  | L. PeEc |  |  |
| L. RSC |  |  | L. FFC | 6 | 3.28 |
| L. STSda | 137 | 4.17 | L. TE2p | 8 | 2.78 |
| L. STSva |  |  | R. 13l | 7 | 2.38 |
| L. A5 |  |  | R. PGs | 12 | 2.44 |
| L. 9p | 38 | 2.91 | R. 8BM | 8 | 2.09 |
| L. 9m |  |  | R. 44 | 38 | 3.56 |
| L. 8Ad |  |  | R. 6r |  |  |
| L. 8Ad | 34 | 3.73 | R. FOP5 | 5 | 2.02 |
| L. 8BL |  |  | R. STSda | 14 | 1.93 |
| L. AIP | 139 | 3.37 | R. STSda | 29 | 3.21 |
| L. PF |  |  | R. STGa |  |  |
| L. IP2 |  |  | R. PFm | 13 | 2.57 |
| L. 47l | 121 | 4.54 | R. PGi | 38 | 2.46 |
| L. 45 |  |  | R. EC | 11 | 1.81 |
| L. IFSa |  |  | R. FFC | 14 | 3.03 |
| L. 46 | 28 | 3.28 | R. TF | 17 | 2.91 |
| L. p9-46v |  |  | R. v23ab | 5 | 2.14 |
| L. PGi | 240 | 4.4 | R. POS1 | 39 | 2.66 |
| L. PGp |  |  | R. RSC |  |  |
| L. PGs |  |  | R. v23ab |  |  |
| L. STSdp | 147 | 4.27 | R. FFC | 9 | 3.84 |
| L. PHT |  |  | R. LO1 | 7 | 1.95 |
| L. STSvp |  |  | R. PIT | 10 | 4.84 |
| L. FFC | 8 | 3.94 | R. PIT | 11 | 4.69 |
| L. FFC | 7 | 3.84 | R. V4 | 13 | 3.66 |
| L. V8 | 8 | 3.18 | R. PIT |  |  |
| L. FFC | 8 | 3.09 | R. V4 | 15 | 3.37 |
| L. V8 | 5 | 3.21 | R. V1 | 117 | 4.12 |
| L. V1 | 239 | 5.36 | R. V2 |  |  |
| L. V4 |  |  | R. V3 |  |  |
| L. V2 |  |  | R. V4 | 9 | 4.73 |
| L. V4 | 18 | 4.07 | R. V3 | 9 | 4.35 |
| L. FFC |  |  | R. V3 | 8 | 4.27 |
| L. V8 |  |  | R. FFC | 8 | 3.59 |
| L. TE2p | 7 | 3.18 | R. PH | 5 | 3.16 |
| L. TE2p | 10 | 3.18 |  |  |  |

**Table S3.**
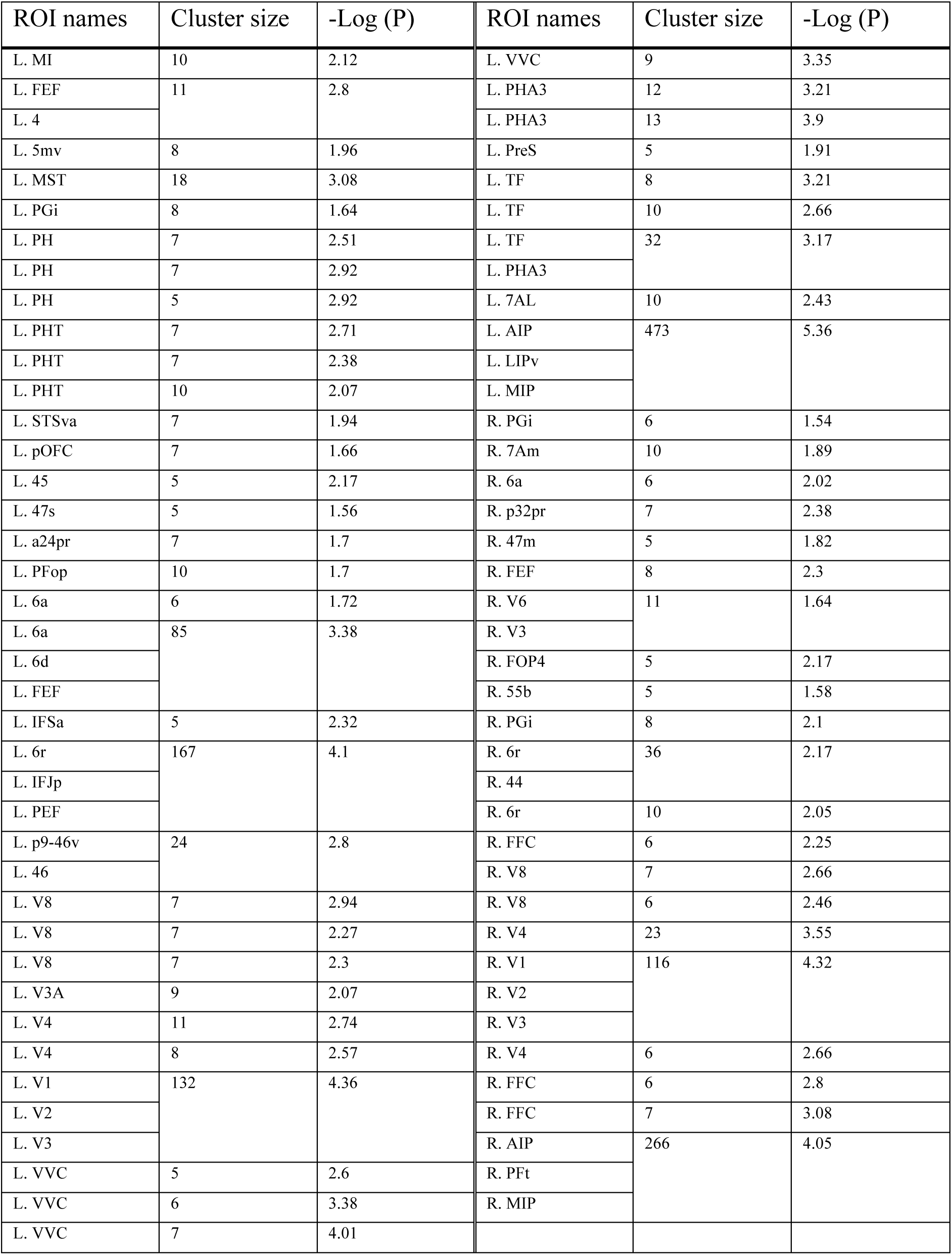
Math-specific brain regions, obtained using Math-to-Math vs. average of Lang-to-Math and Math-to-Lang contrast. Only clusters with at least 5 vertices are shown.

